# *Prochlorococcus* extracellular vesicles: Molecular composition and adsorption to diverse microbes

**DOI:** 10.1101/2020.12.18.423521

**Authors:** Steven J. Biller, Rachel A. Lundeen, Laura R. Hmelo, Kevin W. Becker, Aldo A. Arellano, Keven Dooley, Katherine R. Heal, Laura T. Carlson, Benjamin A. S. Van Mooy, Anitra E. Ingalls, Sallie W. Chisholm

**Author notes:** Corresponding author: Steven J. Biller, Department of Biological Sciences, 106 Central St., Wellesley, MA 02481. Current address: Department of Biological Sciences, Wellesley College, Wellesley, MA.

## Abstract

Extracellular vesicles are small (~50–200 nm diameter) membrane-bound structures released by cells from all domains of life. While vesicles are abundant in the oceans, our understanding of their functions, both for cells themselves and the emergent ecosystem, is in its infancy. To advance this understanding, we analyzed the lipid, protein, and metabolite content of vesicles produced by the marine cyanobacterium *Prochlorococcus*. We show that *Prochlorococcus* exports an enormous array of cellular compounds into the surrounding seawater within vesicles. Vesicles produced by two different strains contain some materials in common, but also display numerous strain-specific differences, reflecting functional complexity within natural vesicle populations. *Prochlorococcus* vesicles contain active enzymes, indicating that they can mediate extracellular biogeochemical reactions in the ocean. We demonstrate that vesicles from *Prochlorococcus* and other bacteria associate with diverse microbes including the most abundant marine bacterium, *Pelagibacter*. Our observations suggest that vesicles may play diverse functional roles in the oceans, including but not limited to mediating energy and nutrient transfers, catalyzing extracellular biochemical reactions, and mitigating toxicity of reactive oxygen species. These findings indicate that a portion of ‘dissolved’ compounds in the oceans are not truly dissolved, but are instead packaged within locally structured, particulate vesicles.

## Introduction

Many, if not all, bacteria release membrane vesicles from their surface into the local environment (Deatherage and Cookson, 2012). In exponentially-growing Gram-negative bacteria these structures derive primarily from the outer membrane, wherein a local region of membrane separates from the cell, carrying with it periplasmic material and other cellular components (Schwechheimer and Kuehn, 2015). A small subset of vesicles from Gram-negative cells may include both outer and inner membrane material as well, further expanding the range of potential vesicle contents (Pérez-Cruz *et al.*, 2015). Vesicles are released constitutively during growth, but release rates can also vary in response to environmental perturbations (MacDonald and Kuehn, 2013; Biller *et al.*, 2014). Extracellular vesicles represent a versatile secretion mechanism for cells (Schwechheimer and Kuehn, 2015; Guerrero-Mandujano *et al.*, 2017), and many classes of cellular compounds, including proteins, nucleic acids, and small molecules, have been identified within bacterial vesicles (Schwechheimer and Kuehn, 2015; Brown *et al.*, 2015). Since they are bounded by a lipid membrane, vesicles may be a particularly useful mechanism for exporting hydrophobic compounds into aqueous extracellular environments (Mashburn-Warren and Whiteley, 2006).

Extracellular vesicles can shuttle their contents between cells (Kadurugamuwa and Beveridge, 1996; Yaron *et al.*, 2000). This ability enables vesicles to mediate a wide variety of biological functions such as horizontal gene transfer, signaling, pathogenesis, quorum signaling, biofilm development, nutrient exchange, viral interactions, and cellular defense (Kadurugamuwa and Beveridge, 1996; Yaron *et al.*, 2000; MacDonald and Kuehn, 2012; Schwechheimer and Kuehn, 2015; Lynch and Alegado, 2017; Schatz *et al.*, 2017). Although such exchanges have been shown to occur both among bacteria and across domains in a few laboratory models, the ‘rules’ dictating these exchanges are not at all clear, and nothing is known about what occurs between cells and vesicles in complex microbial systems such as those found in the oceans, soils, or within the human microbiome.

Extracellular vesicles can reach concentrations of >10^5^ and >10^6^ mL^−1^ in the open ocean and coastal waters, respectively (Biller *et al.*, 2014), and represent an entirely new dimension of dissolved organic carbon pools. Their biological and ecological roles are unknown, but we do know that they are released by diverse taxa of both autotrophic and heterotrophic marine microbes, including the abundant cyanobacterium *Prochlorococcus* (Biller *et al.*, 2014). With a global population of ~3 × 10^27^ cells, *Prochlorococcus* is an important primary producer, responsible for nearly 10% of marine net primary production (Flombaum *et al.*, 2013). This group is known to secrete a number of organic compounds (Bertilsson *et al.*, 2005) that provide marine heterotrophs with a source of carbon and energy (Ottesen *et al.*, 2014; Becker *et al.*, 2019), and contribute to the pool of marine dissolved organic carbon. *Prochlorococcus* extracellular vesicles represent at least one component of this labile organic photosynthate, as purified vesicles have been shown to support the growth of a marine heterotroph (Biller *et al.*, 2014).

Here we explore the contents and function of extracellular vesicles produced by *Prochlorococcus*, focusing on their potential contribution to ocean nutrient pools and ability to interact with other marine microbes. We characterized the lipidome, proteome, and metabolome of vesicles released by two strains, representing high- and low-light adapted ecotypes, and use these inventories to develop hypotheses as to the functions of these structures. To establish whether *Prochlorococcus* vesicles can chemically interact with surrounding seawater, we determined whether they contain active enzymes. Finally, we investigated the potential for *Prochlorococcus* vesicles to mediate biotic interactions by studying whether they can form specific associations with other strains representative of abundant marine microbes.

## Results and Discussion

### What biomolecules are associated with Prochlorococcus extracellular vesicles?

A combination of targeted and untargeted ‘omics’ approaches were used to examine the contents of vesicles and the cells that released them. We focused on vesicles isolated from exponentially-growing, asynchronous cultures of two ecologically distinct *Prochlorococcus* – MIT9312, a high-light adapted strain, and MIT9313, a low-light adapted strain (Biller *et al.*, 2015). We uncovered a vast diversity of biomolecules in these discrete, lipid-bound, colloidal packets, as detailed below.

#### Lipids and pigments

Lipids from vesicles and cells of both *Prochlorococcus* strains were analyzed using an untargeted, high resolution mass spectrometry-based approach. The vesicles contained a suite of different intact polar lipids (IPLs), various pigments, and plastoquinone. Overall, IPLs were relatively more abundant than pigments in vesicles than was seen in the parent cells (Fig. 1a; Table S1). The carotenoids zeaxanthin and carotene were the most abundant pigments in vesicles from both strains, which contained relatively few chloropigments such as divinyl chlorophylls *a* and *b* (Fig. 1b; Table S1). While plastoquinone was identified in the vesicles (Fig. 1b; Table S1) they contained little thylakoid material overall, suggesting a different cellular origin for vesicle carotenoids such as the outer membrane, as has been shown for *Synechocystis* PCC6714 (Jürgens and Weckesser, 1985). Carotenoids have also been observed in vesicles from *Synechocystis* PCC6803 (Pardo *et al.*, 2015), raising broader questions as to whether these compounds might serve a photoprotective role for vesicle contents.

**Figure 1.**
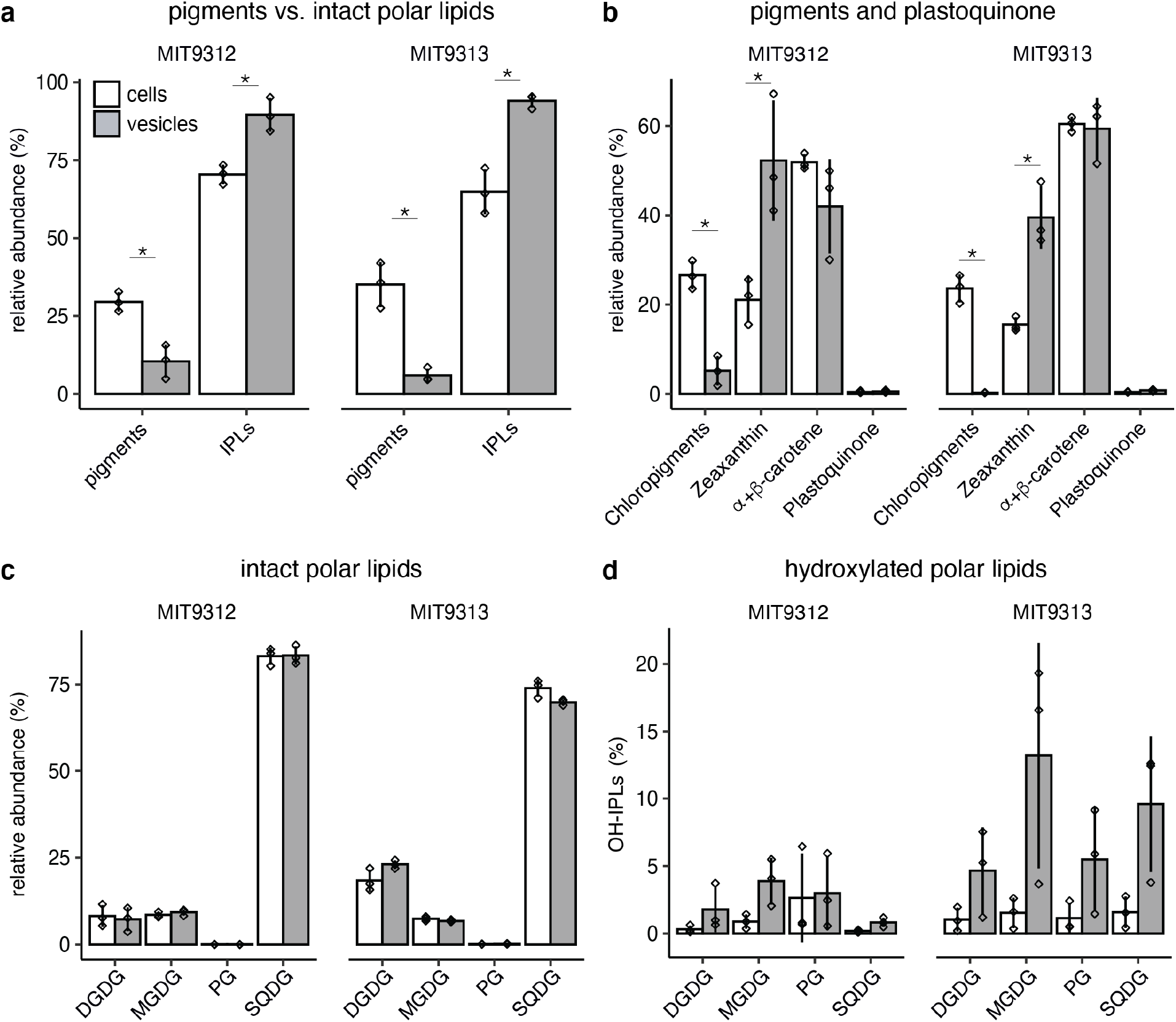
Lipid and pigment content of *Prochlorococcus* cells and vesicles from two strains, MIT9312 and MIT9313. (a) Relative abundance of pigments vs intact polar lipids (IPLs) in vesicles and cells. (b) Relative abundance of specific pigments and plastoquinone. (c) Relative abundance of different intact polar lipid (IPL) groups. (d) Fraction of hydroxylated polar lipids (OH-IPLs) within each IPL class. Values indicate the mean (+/− SD) of three biological replicates of strains MIT9312 (white) and MIT9313 (black). MGDG: monoglycosyl diacylglycerol; DGDG: diglycosyl diacylglycerol; PG: phosphatidylglycerol; SQDG: sulfoquinovosyl diacylglycerol. * indicates significant differences between cells and vesicles (two-tailed *t* test, *p* < 0.05).

The IPL composition of vesicles was dominated by sulfoquinovosyl diacylglycerol (SQDG), diglycosyl diacylglycerol (DGDG) and monoglycosyl diacylglycerol (MGDG) – each found in similar relative abundances (as a proportion of all IPLs) in cells and vesicles (Fig. 1c, Table S1). The overall vesicle lipid composition of each strain more closely reflected the composition of the parent cells than the vesicles from the other strain (Figs. 1c, S1) suggesting that there is not a universal *Prochlorococcus* vesicle lipidome, consistent with our previous observations from *Prochlorococcus* strains MIT9313 and MED4 (Biller *et al.*, 2014). Vesicles from the two strains did share some common features in the fine structure of their polar lipid fatty acids that distinguished them from the parental cells. Of note, many of the vesicle samples had a greater relative abundance of certain oxidized IPLs as compared to cells (Fig. 1d); further, the fraction of some oxidized IPLs (DGDG in MIT9312, SQDG and MGDG in MIT9313) was significantly more variable than in the cells (*F*-test, *p* < 0.05; Figs. 1d, S1b-d). Annotation of these hydroxylated lipids revealed that the hydroxylation is site-specific, for example, at the Δ9-position of a C_18:2_ fatty acid (Fig. S2) regardless of head group. The exact oxidative mechanism (enzymatic, radical-mediated, or non-radical mediated) responsible for generating the hydroxylation is not known, but free radical (auto) oxidation at a specific double bond would be expected to result in at least four isomers of similar abundance (Rontani and Belt, 2020). While lipid hydroxylation could have occurred either before or after vesicle release, the site-specificity raises the possibility that they could be of biological origin. One potential explanation is that *Prochlorococcus* outer membranes are generally enriched in hydroxylated lipids, as has been shown for other Gram-negative bacteria (Volkman *et al.*, 1998). This would be consistent with their enrichment in outer membrane-derived vesicles, and perhaps reflect a role for vesicle secretion as a mechanism for removing damaged compounds from the cell. Hydroxy lipids may also be involved in the membrane curvature and bending processes required for vesicle formation, leading to their preferential incorporation and export.

#### Proteins

Since many of the functional roles for vesicles in other microbial systems are associated with the activity of proteins (Schwechheimer and Kuehn, 2015), we explored the global proteomes of the vesicle and cellular fractions of two *Prochlorococcus* strains using a label-free, quantitative shotgun proteomics approach. During extraction and sample preparation, we utilized surfactants compatible with mass spectrometry to improve the recovery of more membrane-bound proteins and better facilitate in-solution protease digestion, which further helped generate more comprehensive vesicle and cellular proteomes. MIT9312 and MIT9313 vesicles contained, respectively, at least 11% and 12.5% of all predicted proteins encoded by the cell’s genome (Table 1). This most likely does not represent every protein found in vesicles, but rather what could be detected within the relatively small amount of vesicle biomass we could obtain from 20 L cultures as compared to cellular material (c.f., cellular proteomes for MIT9312 and MIT9313 recovered nearly 52% and 37% of all predicted proteins, respectively; Table 1). That said, these vesicle proteomes yielded significantly more protein identifications than our previous study of MIT9313 and MED4 vesicles (in which gel-extracted protein bands were analyzed; Biller *et al.*, 2014), and it is noteworthy that 25 of the 27 proteins we previously identified in MIT9313 were also present in this data set. Further, the 25 were among the top 50 most highly abundant proteins in the MIT9313 vesicle proteome (Tables S2, S3). Notably, the concordance of these results shows that *Prochlorococcus* cells grown under similar conditions will reproducibly package and export intact proteins within vesicles.

**Table 1.** Numbers of proteins identified in *Prochlorococcus* cells and vesicles.

| Strain | Annotated proteins encoded in genomes (Uniprot) | Annotated proteins shared by both genomes | Proteins detected in cells<sup>1</sup> | Proteins detected in cells of both strains | Proteins detected in vesicles | Proteins detected only in vesicles and not in cells | Proteins detected in vesicles from both strains |
| --- | --- | --- | --- | --- | --- | --- | --- |
| MIT9312 | 1964 | 1532 | 1030 | 714 | 224 | 17 | 111 |
| MIT9313 | 2830 |  | 1055 |  | 355 | 41 |  |

<sup>1</sup>These values reflect the data gathered for the trypsin/Lys-C digestion of *Prochlorococcus* cells and vesicles (Tables S2, S3); not included here, additional cellular proteins were identified using Glu-C digestion

Vesicles produced by the two strains had a number of proteins in common (Tables 1, S2), but there were also differences – as has been found in vesicles from other closely related microbes (Tandberg *et al.*, 2016). The vesicle proteomes consisted of proteins associated with many broad functional categories (Figs. 2a, S3), and those most abundant in vesicles from both strains were primarily putative porins (e.g. PMT9312_1131, Som), peptidases/hydrolases (e.g. PMT9312_0677, PMT_1636), chaperones, and uncharacterized proteins (Figs. 2a, S4; Table S4). Proteins uniquely found in MIT9312 vesicles included a cAMP phosphodiesterase (PMT9312_0858), an adhesin-like protein (PMT9312_1179), a phosphate ABC transporter substrate-binding protein (PstB), and some ribosomal proteins. Among those found uniquely in MIT9313 were multiple ABC transporter binding proteins (e.g. UrtA, FutA1, PMT_2203), a sulfatase (PMT_1515), and a putative phosphatase (PMT_1619) (Fig. 2a, Tables S3-S4). Thus, the vesicles released by two relatively similar organisms have distinct functional potentials.

**Figure 2.**
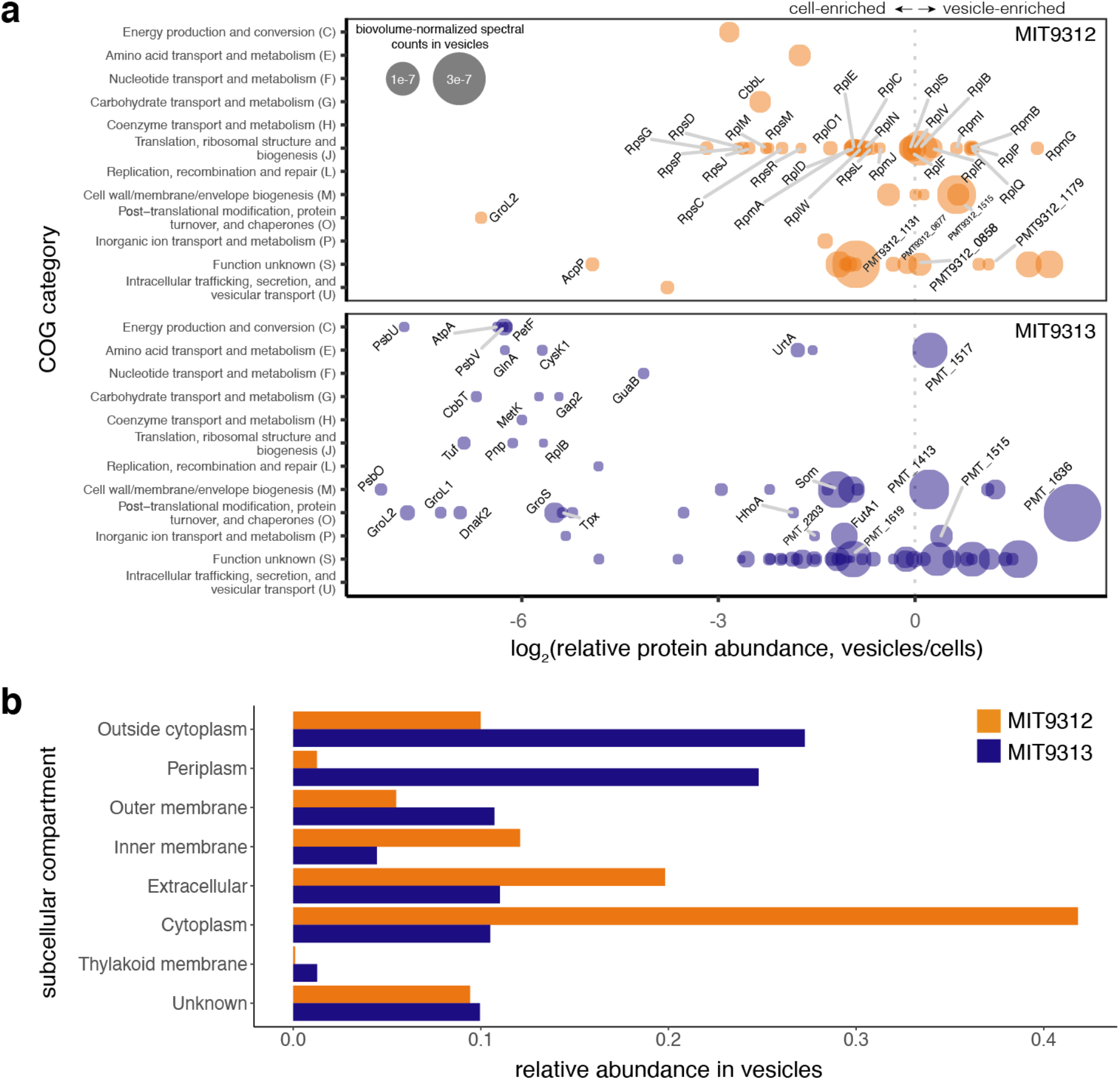
*Prochlorococcus* vesicle proteomes. (a) Relative protein enrichment in *Prochlorococcus* vesicles compared with cells. Points represent the log2 ratio of relative protein abundance for the top 25% most abundant proteins identified in vesicles of strain MIT9312 (above; orange) or MIT9313 (below; blue), as grouped by NCBI clusters of orthologous groups of proteins (COG) functional categories. The area of each point indicates biovolume-normalized spectral counts (abundance) of that protein within the vesicle proteome. Data for all vesicle proteins are found in Table S4. (b) Relative abundance of proteins found in MIT9312 (orange) and MIT9313 (blue) vesicles, based on predicted subcellular localization.

Vesicles are thought to arise primarily from the outer membrane of Gram-negative cells, and indeed we found most of predicted outer membrane and periplasm proteins at some level within vesicles (Fig. 2b, S5, S6a). Studies of other Gram-negative vesicles have identified proteins derived from all cellular compartments (Pérez-Cruz *et al.*, 2015; Zakharzhevskaya *et al.*, 2017; Yun *et al.*, 2017), but we were surprised to find that cytoplasmic proteins made up the largest fraction of vesicle proteins in strain MIT9312 (Fig. 2b); most proteins in the vesicles of the other strain, by contrast, were predicted to originate in the periplasm or an undetermined location outside of the cytoplasm (Fig. 2b). A number of abundant cytoplasmic proteins, such as ribosomal proteins, were found in MIT9312 vesicles (Fig. 2; Tables S2, S3), but there was no general relationship between their abundance in the cells and vesicles (Fig. S6a). Together with the relative lack of thylakoid proteins and chlorophyll in the vesicles, this argues that the presence of cytosolic proteins is likely not due to artifacts such as cell lysis, and that other mechanisms led to their incorporation. Though we cannot rule out the possibility that some individual proteins were specifically packaged via unknown mechanisms, our data support a model wherein proteins are, on average, packaged stochastically into vesicles, with the probability of export influenced by both relative cellular protein abundance and subcellular localization (Fig. S5; S6b-e).

A number of factors likely contribute to the differences observed between the two strains’ vesicle proteomes. Some differences are simply attributable to strain-specific genomic differences (Table 1). While the two genomes share over 1500 genes, MIT9313 encodes ~870 more genes than does MIT9312, and its vesicles contained a proportionally more diverse set of proteins. One notable example was the identification of several prochlorosins, or ProcA peptides, within the vesicle proteome of MIT9313 (Table S2). Prochlorosins are cyclic peptide secondary metabolites encoded by some low-light adapted, but not high-light adapted, *Prochlorococcus* – including 29 diverse *procA* genes in MIT9313 (Li *et al.*, 2010; Cubillos-Ruiz *et al.*, 2017). Strain-specific genome differences do not, however, explain all of the vesicle proteome variations. Even when considering only the proteins shared by both strains, we found that the relative abundance of orthologous proteins was linearly correlated in the whole cell fraction, but that there was no clear relationship in the relative amounts of these proteins within extracellular vesicles (Fig. S3b). This suggests that the packaging of protein cargo into vesicles is either not regulated by the same mechanisms in both strains, or is not actively regulated at all and instead simply reflects underlying variation within cells.

#### Metabolites

A combination of targeted and untargeted metabolomics analysis revealed that vesicle populations from *Prochlorococcus* MIT9312 and MIT9313 contained more than ~1600 and ~2000 unique mass features, respectively, representing molecules with a range of polarities and charge states (Tables S5, S6). Overall, vesicles share the majority of non-polar metabolites with cells, but relatively few polar metabolites are present within vesicles (Tables S5, S6). The general lack of polar molecules in vesicles supports the idea that these structures are effective vehicles for the secretion of hydrophobic compounds (Mashburn and Whiteley, 2005; Schertzer *et al.*, 2009), allowing biological functions that depend on extracellular transport of nonpolar molecules to occur within aquatic ecosystems

Though we are not able to identify most of these mass features, we did observe a number of known compounds within the vesicles (Table 2). For instance, vesicles from both strains contained phylloquinone (Vitamin K1), a compound involved in electron transfer. This result, together with the identification of plastoquinone (Fig. 1b), suggests that vesicles might be involved in dissipating oxidative stress or mediating intercellular electron transfer. Further insights come from the presence of a number of oxidized carotenoid products (Table 2) in vesicles from both strains. Taken together with the previously discussed oxidized IPLs (Fig. 1d), this suggests vesicles could be acting either as a mechanism for removing damaged and unrepairable compounds (Linster *et al.*, 2013), or as a ‘sink’ for reactive oxygen species (ROS) generated within the cell, within vesicles, or in the extracellular seawater environment (Morris *et al.*, 2016). Regardless of the oxidative mechanism(s) involved, suppression of ROS toxicity and/or removal of oxidized metabolites may be another way in which vesicles serve a protective role for cells (Manning and Kuehn, 2011).

**Table 2.** Non-intact polar lipid metabolites identified within *Prochlorococcus* vesicles. For details on analytical fractions and confidence level assessments, see Methods and Supplementary Information. + = detected, nd = not detected.

| Compound name | Fraction | Compound Class | MIT9312 vesicles | MIT9313 vesicles | Confidence level | *m/z* | RT (min) |
| --- | --- | --- | --- | --- | --- | --- | --- |
| Triose | HILICAqNeg | sugar | + | + | 3a | 503.1629 | 13.6 |
| Tetraose | HILICAqNeg | sugar | + | + | 3a | 665.2164 | 14.4 |
| Phylloquinone (Vitamin K1) | RPOrgPos | electron transfer | + | + | 1 | 451.3572 | 14.6 |
| Carotene | RPOrgPos | pigment | + | + | 1 | 536.4371 | 16.1 |
| Protoporphyrin | RPOrgPos | pigment or cyt-c precursor | + | + | 2 | 563.2656 | 12.8 |
| 1-Lauroyl-glycerol (C12) | RPOrgPos | lipid | + | + | 2 | 257.2111 | 11.7 |
| 1-Myristoyl-glycerol (C14) | RPOrgPos | lipid | + | + | 2 | 285.2423 | 13.2 |
| Beta-Apo-12'-carotenal | RPOrgPos | pigment oxidation product | + | + | 3b | 351.2682 | 13.5 |
| Apo-12'-zeaxanthinal | RPOrgPos | pigment oxidation product | + | + | 3b | 367.2633 | 12.0 |
| Beta-Apo-10'-carotenal | RPOrgPos | pigment oxidation product | + | + | 3b | 377.2839 | 13.8 |
| Apo-10'-zeaxanthinal | RPOrgPos | pigment oxidation product | + | + | 3b | 393.2785 | 12.4 |

Though we focused our analysis on the compounds present in vesicles, cellular compounds absent in vesicles are also of interest (Table S7). For example, *Prochlorococcus* cells contained potential osmolytes such as sucrose, glycine betaine, or glucosylglycerol, which help maintain cellular osmotic balance in cyanobacteria (Klähn and Hagemann, 2010), but these compounds were undetectable in either set of vesicles (Table S7). This represents a puzzle as to how cytoplasmic material could be exported to vesicles without including (or retaining) the most abundant solutes in the cell, and how vesicles handle osmotic stresses. Given the diverse roles osmolytes play within cells, such as maintaining enzyme activity and providing redox balance (Yancey, 2005), their absence in the vesicles is intriguing.

### How might vesicles contribute to energy transfer in marine ecosystems?

*Prochlorococcus* vesicles have been shown to serve as sources of organic carbon and/or energy for supporting growth of co-cultured bacteria (Biller *et al.*, 2014). While much of a vesicle’s carbon and chemical energy content is found within the lipids – each 100 nm diameter vesicle will contain ~10,000 lipid molecules in its envelope – proteins and metabolites contribute as well. For example, vesicles from both strains contained triose and tetrose sugars (Table 2), which were not measurable above background levels within cells. These sugars could serve as an additional energy or carbon source for organisms, act as antioxidants, or possibly serve as the primary osmolyte within vesicles (Goh *et al.*, 2010).

Vesicles released from some Gram-negative pathogens have been found to contain ATP (Pérez-Cruz *et al.*, 2015), representing another potential mechanism for vesicle-mediated energy flow. Indeed, we found that vesicles from both strains of *Prochlorococcus* contained ATP. While there was more in MIT9312 than MIT9313 (c.f., 3.6 ± 1.4 × 10^−3^ and 0.13 ± 0.097 × 10^−3^ femtomoles ATP per 10^6^ vesicles, respectively), these amounts represent an average of less than ~2 molecules ATP per vesicle. Although we cannot tell whether this ATP was located in the lumen of the vesicle or associated with some other component (such as a protein), its presence allows for potential vesicle-mediated ATP exchange among cells in the ocean. The presence of proteins and ATP within the same vesicle could facilitate ATP-requiring extracellular reactions as well. This finding has relevance for interpreting measurements of ATP in seawater, where it is found within the ‘dissolved’ (< 0.2μm) fraction – which we now know contains vesicles – and is known to be utilized by marine microbes for biosynthesis (Azam and Hodson, 1977). This dissolved ATP is postulated to come from grazing and/or cellular excretion (Nawrocki and Karl, 1989; Björkman and Karl, 2001). Our results suggest that at least some fraction of ATP in seawater was likely ‘excreted’ via bacterial extracellular vesicles, and is not truly ‘dissolved’.

### Can vesicles transport nutrients?

In an earlier study we observed nutrient binding proteins, such as those for urea, phosphate, and iron, in the vesicles exported by *Prochlorococcus* strains MED4 and MIT9313 (Biller *et al.*, 2014). This suggested that vesicles might transport nutrients or even scavenge compounds as they diffuse through seawater and, in turn, organisms that encounter these vesicles could gain access to a locally concentrated ‘packet’ of nutrients. To explore this idea further, we looked for transporters and binding proteins that could facilitate the transfer of either organic or inorganic compounds in our proteome data. Vesicles produced by both strains contained putative transporters for sulfate, magnesium, and ammonium (Tables S2, S3) suggesting that – assuming favorable energetics and proper protein orientation in the membrane – vesicles could concentrate these compounds. We also identified numerous putative substrate-binding proteins (many associated with ABC transporters) for iron, manganese, urea, amino acids, and phosphate in the vesicles (Tables S2, S3) that could serve as a mechanism through which vesicles bind these compounds, thus avoiding losses via diffusion. To explore this possibility, we measured vesicle phosphate content and compared it with that of the average cell. Vesicles from MIT9312 and MIT9313 contained 3.4 ± 3.2 and 3.7 ± 2.1 femtomoles phosphate per 10^6^ vesicles, respectively. Given that a *Prochlorococcus* MED4 cell (similar in size to MIT9312) contains ~10 attomoles phosphorus (P) (Bertilsson *et al.*, 2003) and *Pelagibacter* HTCC7211 cells contain ~16 attomoles P (White *et al.*, 2019) under P-limited conditions, an encounter with a single *Prochlorococcus* vesicle would supply only ~0.01% of these cells’ P quota. These are averages, however, and absent an understanding of the distribution of P per vesicle, and vesicle-cell encounter rates, it is impossible to speculate further about whether vesicles may serve as a significant source of inorganic nutrients on a cell-by-cell basis in the oceans. Finally, we found putative ssDNA, dsDNA, and RNA-binding proteins in vesicles (Tables S2, S3), and note that vesicle-associated nucleic acids may also function as a nutrient source for microbes (Jørgensen and Jacobsen, 1996).

### Do Prochlorococcus vesicles have enzymatic activity?

Many of the most abundant proteins in the vesicles were putative enzymes (Table S2) with the potential to mediate exoenzymatic activities. Indeed, vesicles from both *Prochlorococcus* strains exhibited active lipase, phosphatase, protease, and sulfatase activities (Fig. 3), indicating that they could contribute to biogeochemical cycling *in situ.* Vesicles thus provide a mechanism for cells to catalyze extracellular reactions, potentially providing them access to extracellular substrates they might not otherwise encounter (Ebner and Götz, 2019). For instance, we found that vesicles from MIT9313 displayed measurable sulfatase activity while intact cells did not (Fig. 3d). Further, enzymes enclosed within vesicles could be protected from environmental damage and degradation, allow for simultaneous co-secretion of multiple proteins, and/or provide an environment where substrates could be maintained in close proximity in the otherwise dilute oligotrophic ocean (Bonnington and Kuehn, 2014). While cyanobacteria and many other microbes are known to specifically secrete certain exoenzymes (Christie-Oleza *et al.*, 2015), the broad diversity of vesicle-associated enzymes seen here – including those typically considered cytoplasmic – blurs the distinction between intracellular and extracellular enzymes and expands the ways marine microbes could be interacting with the abiotic environment and influencing ocean biogeochemistry.

**Figure 3.**
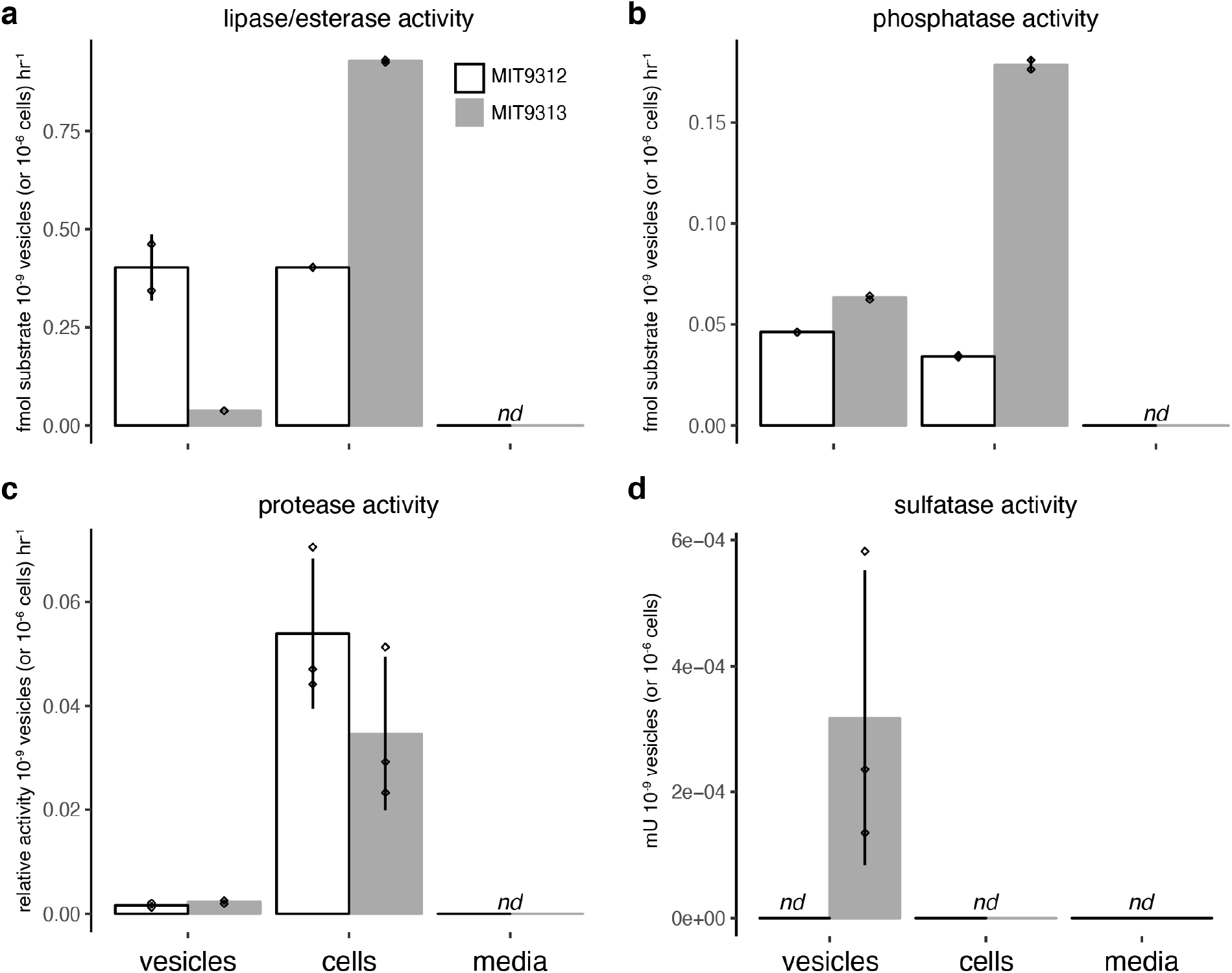
Enzymatic activity in purified *Prochlorococcus* vesicles compared to that of cells and media background. Activity of (a) lipases, (b) phosphatases, (c) proteases, and (d) sulfatases in samples from *Prochlorococcus* MIT9312 (white) and MIT9313 (gray). Values represent the mean (+/− SD) of 2-3 biological replicates, standardized per population of either 10^9^ vesicles or 10^6^ cells. *nd* = not detected.

### Can marine extracellular vesicles interact with cells from different species?

We know from studies of terrestrial and host-associated microbes that many biological functions of vesicles are attributable to the transfer of vesicle contents between cells (Schwechheimer and Kuehn, 2015; Brown *et al.*, 2015). Vesicle-mediated biotic ‘interactions’ may encompass a range of mechanisms, from vesicle-associated activities occurring in proximity to a cell (e.g. enzymatic activities or phage defense; MacDonald and Kuehn, 2012; Biller *et al.*, 2014; Rakoff-Nahoum *et al.*, 2014) to the delivery of vesicle contents into another cell (Kadurugamuwa and Beveridge, 1999). To begin to explore interactions between *Prochlorococcus* vesicles and cells, we looked for physical associations between fluorescently-labeled vesicles and six different marine microbes from the Proteobacteria, Cyanobacteria, and Bacteroidetes – the three most abundant bacterial phyla in the oceans (Fig. 4a). We found that vesicles from *Prochlorococcus* MIT9313 associated with cells of both *Prochlorococcus* MIT9312 and MIT9313, along with representatives of the marine Gammaproteobacteria and Alphaproteobacteria – including *Candidatus Pelagibacter ubique* HTCC7211, a member of the numerically dominant SAR11 group of marine heterotrophs (Figs. 4, S7). The breadth of interactions was not specific to *Prochlorococcus* vesicles, as vesicles purified from the marine heterotroph *Alteromonas* MIT1002, which was co-isolated with *Prochlorococcus* (Biller, Coe, *et al.*, 2015), were also able to interact with these strains (Figs. 4, S7). To ensure that the observed vesicle-cell associations were not an artifact of the fluorescent dye used, we incubated the same set of marine microbes with vesicles from an *E. coli* strain (Dinh and Bernhardt, 2011) which instead expressed GFP within vesicles, and found that they were also able to interact with multiple microbes (Figs. 4b, S7). While we further verified that our fluorescent vesicle labeling approach did not significantly influence the ability of *E. coli* vesicles to interact with cells (Fig. S8), we cannot rule out a potential influence of vesicle surface modifications on the pairwise interactions tested.

**Figure 4.**
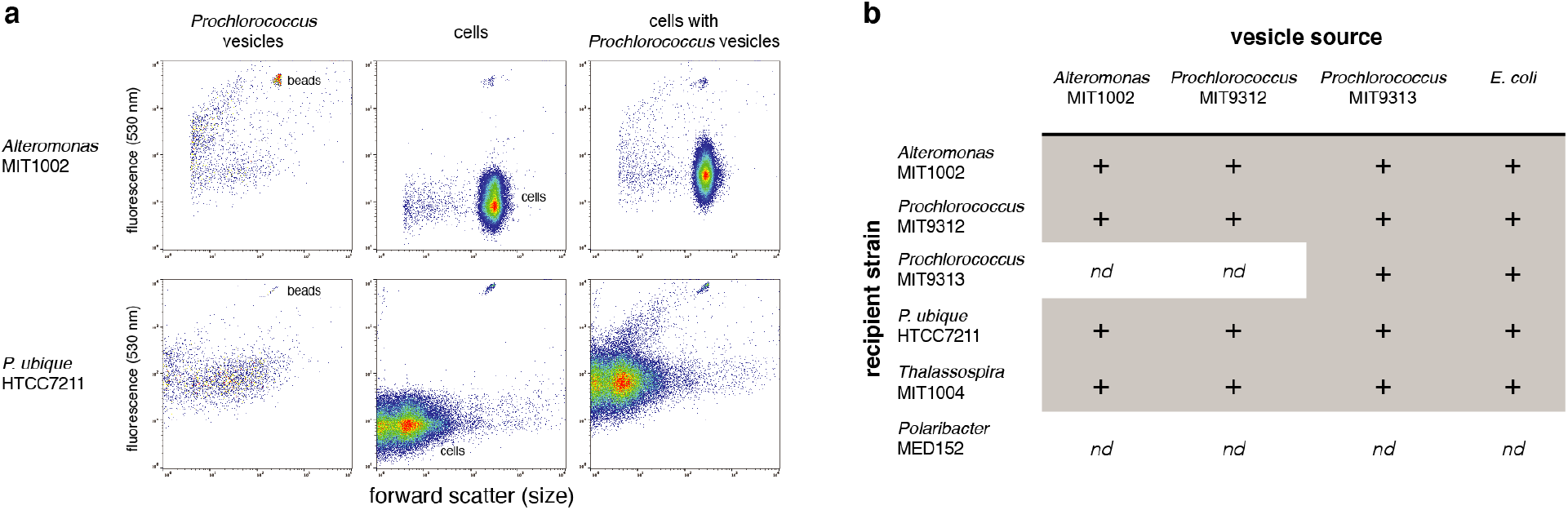
Association of extracellular vesicles with diverse microbial cells. (a) Example flow cytometry plots of fluorescently labeled *Prochlorococcus* MIT9313 vesicles (left), *Alteromonas* MIT1002 and *Pelagibacter ubique* (SAR11) cell populations (center), and cells following a two hour incubation with vesicles (right). Internal reference beads are labeled. (b) Interaction of labeled vesicles with cells (+), or lack thereof (*nd* = not detected), for different ‘source’ strains (vesicle producers) and ‘recipient’ strains (cells exposed to labeled vesicles). Positive interactions were determined based on two criteria: an increase in normalized median cellular 530 nm fluorescence when mixed with labeled vesicles (as in a; see also Fig. S7), and a statistically significant shift in the cellular population distribution (Chi Squared T(x) test, *p* << 0.01 in 2-3 biological replicates; see Supplementary Methods). *Alteromonas* and *Prochlorococcus* vesicles were covalently labeled with an amine-reactive Alexa 488 dye; *E. coli* vesicles contained GFP.

Though four of the strains examined interacted with all of the vesicles tested (Fig. 4), two others exhibited apparent specificity toward vesicles from different sources. That is, vesicles from both *Alteromonas* and *Prochlorococcus* MIT9312 did not associate with *Prochlorococcus* MIT9313 cells, and none of the vesicles interacted with *Polaribacter* MED152, a marine *Bacteroidetes* (González *et al.*, 2008), at our detection levels (Fig. 4b). This confirms that our observations are not simply due to experimental conditions or random ‘sticking’ of vesicles to cells. Specificity in vesicle-cell associations has been shown in enterobacteria (Tashiro *et al.*, 2017), and likely underlies variation in vesicle-mediated DNA transfer rates among species (Tran and Boedicker, 2017). Our data raise the possibility that different vesicle-cell pairs may vary in the strength of their interactions (Fig. S7), for example, the propensity of a vesicle to associate with a particular cell may depend on its specific surface composition, which would likely be obscured by the averaging effect of our population-scale analysis. The apparent lack of vesicle associations with *Polaribacter* in our experiments is striking and invites an exploration of what factors, such as surface charge (Tashiro *et al.*, 2017), cell envelope structure, and hydrophobicity (MacDonald and Beveridge, 2002), might define interaction boundaries.

The vesicle-cell associations could be disrupted by repeated rounds of centrifugation and washing of the cells (Fig. S9), suggesting that the majority of labeled vesicle material was not fully integrated into the outer membrane of the cells (at least at this timepoint), but was attached via non-covalent bonding between the two surfaces. Though these observations do not directly establish a particular biologically-relevant function, we speculate that ‘captured’ vesicles could serve many functions, including delivering cargo through membrane fusion, ‘flipping’ of molecules from the vesicle membrane into the cell (Remis *et al.*, 2014), or extracellular degradation and subsequent uptake of vesicle contents. Alternatively, the presence of vesicles nearby could serve defensive roles for cells (Manning and Kuehn, 2011) or influence cellular processes via enzymatic activities.

### Conclusions

We have shown that vesicles released by *Prochlorococcus* transport diverse cellular components, with the potential to impact both the abiotic and biotic components of marine ecosystems. Our data detail compounds that are secreted and potentially trafficked via *Prochlorococcus* extracellular vesicles, including organic molecules and inorganic nutrients – many of which likely contribute to the flow of energy and nutrients among cells. The metabolomic data further reinforce the potential importance of vesicles as mechanisms for transporting nonpolar compounds in aqueous environments. In addition, the ability of vesicles to transport a diverse suite of active enzymes outside of *Prochlorococcus* cells suggests that a broad array of cellular enzymes could actually be considered exoenzymes, and implicates vesicles as a component of *Prochlorococcus*’ global biogeochemical footprint.

The finding that vesicles from *Prochlorococcus* and other marine bacteria can associate – with some specificity – with cells of other species provides a new route for matter, energy and information exchange among and between microbes in the oceans. Of particular interest is that vesicles from *Prochlorococcus*, the numerically dominant phytoplankter in many regions, can associate with *Pelagibacter*, the most abundant heterotrophic microbe in the oceans. The fact that *Polaribacter* did not show a significant association with any vesicles indicates that there is selectivity in the system, influencing how we might think about vesicles in the context of ‘public goods’ in the oceans.

While the diversity of compounds we found associated with *Prochlorococcus* vesicles already reveals them to be a remarkably versatile cellular secretion system (Guerrero-Mandujano *et al.*, 2017), it is likely that additional sampling would reveal an even longer ‘tail’ of other proteins and metabolites that are either exported at relatively lower abundances or were perhaps lost through the purification protocols used here. Thus we suspect that most (or perhaps all) proteins expressed within cells will eventually be found within vesicle populations. Further, vesicle compositions will undoubtedly vary to some degree with changes in growth conditions (Orench-Rivera and Kuehn, 2016). Given the large diversity of proteins and small molecules identified in the vesicles, it seems implausible that these materials are uniformly distributed among each member of the vesicle population. We have previously shown heterogeneity of DNA fragment incorporation into vesicles from multiple marine bacteria (Biller *et al.*, 2017), and vesicle subpopulations containing both outer and inner membrane material from Gram-negative cells have been reported (Pérez-Cruz *et al.*, 2015). Furthermore, distinct vesicle subpopulations with unique origins and epitopes have been noted for some time in eukaryotic cells (Kowal *et al.*, 2016). Thus the functional potential of one vesicle is likely different from the next – similar to our perspective on the cells from which vesicles are derived, whose ecological role can only be understood by viewing them as populations of diverse entities which may display a diverse range of functions in different contexts (Biller, Berube, *et al.*, 2015).

The abundance of vesicles in seawater, and the diversity of their chemical contents, prompt a reassessment of how we conceptualize and interpret the pool of dissolved organic carbon that is so critical to the function of marine ecosystems. While many cellular compounds are secreted in the form of truly ‘dissolved’ individual molecules, it is now clear that microbes also release organic molecules in locally structured, colloidal vesicles which may influence the accessibility, activity, and half-lives of their contents. As such, vesicles blur the boundaries between definitions of ‘particulate’ and ‘dissolved’, or ‘living’ and ‘detrital’, in marine biogeochemistry – distinctions that are critical for understanding and modeling the system. Extracellular vesicles represent a new frontier for understanding the flow of energy, materials, and information in ocean ecosystems.

## Experimental Procedures

### Culturing conditions and sampling for lipidomics, proteomics, and metabolomics

Axenic cultures of *Prochlorococcus* MIT9312 and MIT9313 used for lipidomics, proteomics, and metabolomics analyses were grown in defined artificial AMP1 media (Moore *et al.*, 2007) supplemented with 10 mM (final concentration) filter-sterilized sodium bicarbonate. All 20 L cultures were grown in polycarbonate carboys (ThermoFisher Nalgene, Waltham, MA, USA) with gentle stirring (60 rpm), under constant light flux (10 – 20 μmol Q m^−2^ s^−1^ for MIT9313; 30 – 40 μmol Q m^−2^ s^−1^ for MIT9312) at 24°C.

All cell and vesicle samples were collected during mid-to-late exponential growth phase. Cell pellets were obtained by gently centrifuging cells (7,500 xg) for 10 minutes at 4°C; vesicle isolation is detailed below. For lipidomics, proteomics, and metabolomics analyses, a total of seven 20 L cultures were grown in AMP1 media for each of the two *Prochlorococcus* strains, providing three replicates for the lipid and small metabolite analysis and an additional sample for proteomics analysis. *Prochlorococcus* AMP1 growth media blanks and vesicle suspension buffer (phosphate buffered saline, PBS) blanks served as controls for lipidomics, proteomics, and metabolomics analysis; they were extracted and analyzed alongside all cell and vesicle samples. Since *Prochlorococcus* was grown under continuous light, the cultures were not synchronized. Thus, all cell samples represent a bulk average population of cells at all stages of the cell cycle; vesicle samples similarly integrate material released across both the lag and exponential growth (steady-state) phases.

### Vesicle isolation

Vesicles were isolated as described previously (Biller *et al.*, 2014). Briefly, cultures were first gravity filtered through a 0.2 μm capsule filter (Polycap 150TC; GE Life Sciences/Whatman, Maidstone, UK). The filtrate was then concentrated using a 100 kDa tangential flow filter (Ultrasette with Omega membrane; Pall, Port Washington, NY, USA) and re-filtered through a 0.2 μm syringe filter. Vesicles were pelleted from the sample by ultracentrifugation at ~100,000 x g (Beckman-coulter SW32Ti rotor; 32,000 rpm, 1.5 hrs, 4°C), purified on an OptiPrep gradient (Biller *et al.*, 2014), then washed and resuspended in 0.2 μm filtered 1x PBS.

Vesicle concentrations were measured with a NanoSight LM10HS instrument equipped with a LM14 blue laser module using NTA software V3.1 (NanoSight / Malvern, Westborough, MA, USA). Samples were diluted such that the average number of particles per field was between 20-60, per the manufacturer’s guidelines. Three replicate videos were collected from each sample at a camera level of 11, and analyzed at a detection threshold of 1. The sample chamber was thoroughly flushed with 18.2 MΩ cm^−1^ water (Milli-Q; Millipore, Burlington, MA, USA) between samples, and visually examined to ensure that no particles were carried over.

### Lipidomics

Lipids were extracted from triplicate cell pellets and vesicles from *Prochlorococcus* MIT9312 or MIT9313 using a modified Bligh and Dyer protocol (Popendorf *et al.*, 2013). The total lipid extract was analyzed on an Agilent 1200 reversed phase high performance liquid chromatograph (HPLC) coupled to a ThermoFisher Exactive Plus Orbitrap high resolution mass spectrometer (HRMS; ThermoFisher, Waltham, MA, USA). HPLC and MS conditions are described in Collins *et al.*, 2016 (modified after Hummel *et al.*, 2011). For the identification and peak area integration, we used LOBSTAHS, an open-source lipidomics software workflow based on adduct ion abundances and several other orthogonal criteria (Collins *et al.*, 2016). Lipid peaks identified using the LOBSTAHS software were integrated from MS data after pre-processing with XCMS (Smith *et al.*, 2006) and CAMERA (Kuhl *et al.*, 2012) and corrected for differences in response factors among different lipids using commercially available standards as described in (Becker *et al.*, 2018). Additional details are available in Supplementary Information.

### Proteomics

Proteins were extracted from cell biomass and vesicles from a single 20 L batch culture of *Prochlorococcus* MIT9313 or MIT9312. Filters containing cell biomass or vesicle suspensions were extracted using bead-beating and freeze-thaw cycles. Proteins from lysed cells or vesicles were suspended in RapiGest SF (Waters, Milford, MA, USA) to facilitate protein solubilization and then underwent disulfide reduction with tris(2-carboxyethyl)phosphine, alkylation with iodoacetamide, and in-solution protease digestion using either a trypsin/Lys-C mix or Glu-C (Promega, Madison, WI, USA). Following RapiGest hydrolysis and desalting, samples were resuspended in a solution containing an internal standard of synthetic peptides (Hi3 *Escherichia coli* standard, Waters), spiked with iRT retention time standard (Biognosys, Boston, MA, USA), and immediately analyzed on a Waters ACQUITY M-class LC coupled to a Thermo QExactive HF HRMS equipped with a nano-electrospray source. Data dependent acquisition was performed on the top 10 ions and data analysis were conducted using the software from the trans-proteomic pipeline (TPP v.5.1.0) (Nesvizhskii *et al.*, 2007).

Label-free comparison of relative protein abundances was facilitated by normalizing protein spectral counts to the Hi3 internal standard to account for differences in ionization of peptides due to sample matrix affects and then normalizing to the amount of biomass injected onto the LC-HRMS. To compare protein enrichments within *Prochlorococcus* cell and vesicle fractions, “per cell” or “per vesicle” spectral counts were then normalized on the basis of size, or biovolume, differences between a cell and a vesicle, herein referred to as “biovolume-normalized protein spectral counts” (Tables S2-S3). Unless otherwise indicated, proteome comparisons across cells and vesicles were conducted using biovolume-normalized spectral counts from only trypsin/Lys-C-based proteomes (Fig. 2). Relative protein enrichments in *Prochlorococcus* vesicles as compared with cells was computed using log_2_ ratios (Table S4) and displayed in Figure 2a using only the proteins with spectral counts in the top 25% of those identified in vesicles of strain MIT9312 or MIT9313. Additional details are available in Supplementary Information.

### Metabolomics

Cell pellets and vesicles originating from triplicate 20 L batch cultures of *Prochlorococcus* MIT9312 or MIT9313 were extracted using a modified Bligh and Dyer technique (Bligh and Dyer, 1959; Boysen *et al.*, 2018). Cell pellets were manually disrupted by bead beating during the extraction, as described by Boysen *et al.*, 2018; vesicles were extracted without bead beating. Metabolite separations were achieved using reversed-phase (RP, for aqueous and organic extracts) and hydrophilic interaction liquid chromatography (HILIC, for aqueous extracts only), as detailed in Table S8 and Boysen *et al.*, 2018. Data were collected in positive ion mode for RP analyses and in positive and negative mode (using polarity switching) for HILIC analyses. For every sample, data were processed in four subgroups, defined by phase, chromatography, and ion mode: RP-organic-positive (RPOrgPos), RP-aqueous-positive (RPAqPos), HILIC-aqueous-positive (HILICAqPos), HILIC-aqueous-negative (HILICAqNeg). Data were collected using Thermo QExactive HF HRMS using same settings as in Boysen *et al.*, 2018.

For targeted data, individual metabolite features were integrated using Skyline Daily (MacLean *et al.*, 2010) and subjected to an in-house quality control protocol (Boysen *et al.*, 2018). Untargeted data were processed using MS-DIAL software (Tsugawa *et al.*, 2015) using parameters reported in Table S9. MS-DIAL was used to pick, align, and integrate mass features from raw datasets. Identification of mass features within untargeted data sets was accomplished via dereplication of the mass feature list against several databases, yielding identifications of variable confidence as described in Heal *et al* 2019. We ranked confidence in mass feature identifications according to existing literature (Sumner *et al.*, 2007). All mass features were searched against an Ingalls Lab in-house database of authenticated standards, LOBSTAHs output, and MassBank (Horai *et al.*, 2010). Additional details are available in Supplementary Information.

### Vesicle enzyme activity assays

All nutrient and enzymatic assays were carried out using vesicles collected from 20 L exponentially growing *Prochlorococcus* cultures grown as above. Vesicles were purified using an OptiPrep (Iodixanol) density gradient as described in Biller *et al.*, 2014, and washed in clean 1x PBS. ATP was measured using a standard luminescence-based assay with the BacTiter-Glo kit (Promega) according to the manufacturer’s directions. Vesicle phosphate was measured using the Sigma-Aldrich Phosphate Assay kit (MilliporeSigma, St. Louis, MO, USA). Protease measurements were made using the Sigma-Aldrich Protease Fluorescent Detection Kit following the manufacturer’s instructions with the following modifications: 10 μl of a 2x incubation buffer was used in each 50 μl reaction, which allowed us to add 20 μl of sample to each reaction and increase the number of vesicles included in the reaction. The reaction was carried out for 18-20 hours at 28 °C in the dark. Sulfatase activity was measured using the BioVision Sulfatase Activity Assay (BioVision, Milpitas, CA, USA), following manufacturer’s instructions.

Phosphatase and lipase activity were measured by following the hydrolysis of fluorescent substrates 4-Methylumbelliferyl phosphate and 4-Methylumbelliferyl oleate (both from Sigma-Aldrich) following standard methods (Hoppe, 1993). Briefly, ~10^10^ vesicles in 1x PBS were incubated with 100 μM substrate (dissolved in ethylene glycol monomethyl ether) and natural seawater from the Sargasso Sea in a 200 μL reaction within a black-walled 96 well plate at room temperature for 16 hours. Boiled vesicle samples and substrate in seawater lacking any added vesicles were used as controls.

### Vesicle interaction experiments

Vesicles were purified from 20 L cultures of *Prochlorococcus* MIT9312 or MIT9313 (grown as above but in natural-seawater based Pro99 media (Moore *et al.*, 2007) containing 0.2 μm filtered water from Vineyard Sound, MA); 10 L cultures of *Alteromonas* strain MIT1002 (Biller, Coe, *et al.*, 2015) grown at 24 °C in ProMM media (Pro99 media, as above, plus lactate, pyruvate, glycerol, acetate, and Va vitamins (Berube *et al.*, 2015)); or 10 L cultures of *E. coli* strain TB28(*att*HKTB263) (Dinh and Bernhardt, 2011; this strain expresses the Superfolder variant of GFP and targets it to the periplasm) grown at 30 °C in M9 maltose media supplemented with 250 μM isopropyl β-D-1-thiogalactopyranoside and 100 μg/mL ampicillin. Purified vesicles were covalently labelled with an amine-reactive Alexa Fluor 488 5-SDP ester dye (Molecular Probes/ThermoScientific).

To examine the ability of vesicles to associate with representative phylogenetically distinct marine microbes, cells of *Prochlorococcus* MIT9312 and MIT9313 (in Pro99 media), *Alteromonas* MIT1002 and *Thalassospira* MIT1004 (Biller *et al.*, 2017) (in ProMM media), and *Polaribacter* MED152 (González *et al.*, 2008) (in Pro99 media, supplemented with 5 g/L peptone and 1g/L yeast extract) were all grown to mid-exponential growth phase at 24 °C. *Pelagibacter* HTCC7211 was grown in AMS1 media at 22 °C. Approximately 10^9^-10^10^ vesicles (labeled or unlabeled), or an equivalent volume of the PBS Alexa 488 labeling control, were added to 2 mL of culture at ~10^5^ cells mL^−1^ (final ratio: ~1000:1 vesicles:cells). Cultures with vesicles were incubated for 2 hours at the normal growth temperatures and examined on an Influx flow cytometer (Cytopeia/BD, Franklin Lakes, NJ). Cells were excited using a blue 488 nm laser and monitored for chlorophyll (692/40 nm emission) and Alexa 488/GFP (530/40 nm emission) fluorescence. All flow cytometry data was analyzed using FlowJo (V10.5). Additional details are available in Supplementary Information.

### Misc

All data were analyzed in R (V3.5.1). Plots were created using ggplot2 (Wickham, 2009).

### Data availability

The mass spectrometry proteomics data have been deposited to the ProteomeXchange Consortium via the PRIDE (Perez-Riverol *et al.*, 2019) partner repository with the dataset identifier PXD013602. Metabolite data are available at the NIH Common Fund's National Metabolomics Data Repository (NMDR) website, the Metabolomics Workbench, https://www.metabolomicsworkbench.org (Sud *et al.*, 2016), under Study ID # ST001524.

## Acknowledgments

This work was funded by grants from the National Science Foundation (OCE-1356460 to S.W.C.) and the Simons Foundation (SCOPE Award ID 329108 to B.A.S.V.M., A.E.I., S.W.C.; Simons Award ID 385428 to A.E.I. and 598819 to K.R.H.). K.W.B was supported by the Postdoctoral Scholarship Program at the Woods Hole Oceanographic Institution. R.A.L was partially supported by a postdoctoral fellowship from the Swiss National Science Foundation. We thank Thomas Bernhardt (Harvard Medical School) for providing the sfGFP-expressing *E. coli* strain, Jarone Pinhassi (Linnaeus U.) for sharing *Polaribacter* MED152, and Stephen Giovannoni (U. Oregon) for *Pelagibacter* HTCC7211.

