## Supplementary Information for "*Prochlorococcus* extracellular vesicles: Molecular composition and adsorption to diverse microbes"

**Supplementary Methods**

*Lipidomics*

Lipids were extracted from cell pellets (approximately 1.8 x $10^{11}$ – 1.7 x $10^{12}$ cells for MIT9312, 2.9 x $10^{11}$ – 5.4 x $10^{11}$ cells for MIT9313) and vesicles (approximately 3.3 x $10^{10}$ – 6.4 x $10^{11}$ for MIT9312, 1.2 x $10^{12}$ – 2.9 x $10^{12}$ for MIT9313) using a modified Bligh and Dyer protocol (Popendorf *et al.*, 2013) with DNP-PE-$C_{16:0}$/$C_{16:0}$-DAG (2,4-dinitrophenyl phosphatidylethanolamine diacylglycerol; Avanti Polar Lipids, Inc., Alabaster, AL, USA) used as an internal recovery standard. Filter blanks, *Prochlorococcus* growth blanks, and vesicle suspension buffer blanks were extracted and analyzed alongside cell and vesicle samples. The total lipid extract was analyzed by reversed phase high performance liquid chromatography (HPLC) mass spectrometry (MS) on an Agilent 1200 HPLC coupled to a ThermoFisher Exactive Plus Orbitrap high resolution mass spectrometer (ThermoFisher Scientific, Waltham, MA, USA). HPLC and MS conditions are described in Collins *et al.*, 2016 (modified after Hummel *et al.*, 2011). In brief: 20 µL were injected onto a C8 Xbridge HPLC column (particle size 5 µm, length 150 mm, width 2.1 mm; Waters Corp., Milford, MA, USA). Lipids were eluted at a flow rate of 0.4 mL $min^{-1}$ using the following gradient with eluent A (water with 1% 1M ammonium acetate and 0.1% acetic acid) and eluent B (70% acetonitrile, 30% isopropanol with 1% 1M ammonium acetate and 0.1% acetic acid): 45% A was held for 1 min, from 45% A to 35% A in 4 min, from 25% A to 11% A in 8 min, from 11% A to 1% A in 3 min with an isocratic hold until 30 min. Finally, the column was equilibrated with 45% A for 10 min. ESI source settings were: Spray voltage, 4.5 kV (+), 3.0 kV (-); capillary temperature, 150 ºC; sheath gas and auxiliary gas, both 21 (arbitrary units); heated ESI probe temperature, 350 °C. Mass data were collected in full scan while alternating between positive and negative ion modes. For each MS full scan, up to three $MS^2$ experiments targeted the most abundant ions with $N_2$ as collision gas. The scan range for all modes was 100-1500 *m/z*. The mass spectrometer was set to a resolving power of 140,000 (FWHM at *m/z* 200) leading to an observed resolution of 75,100 at *m/z* 875.5505 of our internal standard, DNP-PE. Exact mass calibration was performed by weekly infusing a tune mixture. Additionally, every spectrum was corrected using a lock mass, providing real-time calibrations.

For the identification and quantification of lipids, we used LOBSTAHS, an open-source lipidomics software pipeline based on adduct ion abundances and several other orthogonal criteria (Collins *et al.*, 2016). Lipids identified using the LOBSTAHS software were quantified from MS data after pre-processing with XCMS (Smith *et al.*, 2006) and CAMERA (Kuhl *et al.*, 2012). XCMS peak detection was validated by manual identification using retention time as well as accurate molecular mass and isotope pattern matching of proposed sum formulas in full-scan mode and tandem MS ($MS^2$) fragment spectra of representative compounds (Becker *et al.*, 2018). Low abundance compounds were excluded by using a signal to noise ratio cutoff of 10 and defining other peak picking parameters, such as peak width and peak shape, as described by (Collins *et al.*, 2016). As a means of validating the accuracy and reliability of LOBSTAHS identification and quantification, quality control (QC) samples of known composition were interspersed with samples from the *Prochlorococcus* cultures and vesicles. The QC samples contained a mixture of authentic, commercially available lipids that has been used extensively in other work (Van Mooy and Fredricks, 2010; Popendorf *et al.*, 2013; Collins *et al.*, 2016; Becker *et al.*, 2018). Lipid peaks obtained from LOBSTAHS were corrected for the relative response of the same standards that were added to the QC samples. Individual response factors were determined from external standard curves by triplicate injection of a series of standard solution mixtures in amounts ranging from 0.15 to 40 pmol on column per standard. Our method's use of external standards was recently validated in a study that compared lipid quantitation against internal, isotope-labeled standards (Becker *et al.*, 2018). Data were corrected for differences in extraction efficiency using the recovery of the DNP-PE internal standard. The average analytical error for the standards in the QC samples was ± 6.1%. Standards for phosphatidylglycerol (PG) and DNP-PE were purchased from Avanti Polar Lipids, Inc., (Alabaster, AL, USA). Purified natural MGDG and DGDG were purchased from Matreya LLC (Pleasant Gap, PA, USA) or Sigma-Aldrich (St. Louis, MO, USA), and purified sulfoquinovosyl diacylglycerol (SQDG) from spinach was purchased from Lipid Products (South Nutfield, UK). Response factors for plastoquinones were determined using a ubiquinone ($UQ_{10:10}$) standard, chlorophylls and pheophytins using a chlorophyll *a* standard, and zeaxanthin and $\alpha/\beta$-carotene using a $\beta$-carotene standard all purchased from Sigma Aldrich.

*Proteomics*

Vesicles were collected from single 20 L batch cultures of MIT9313 or MIT9312 and gradient purified, then washed three times in phosphate-buffered saline (PBS). A control sample from 20 L of AMP1 artificial seawater media only was also collected. Vesicle abundances corresponded to $1.1 \times 10^{12}$ particles/mL and $7.3 \times 10^{11}$ particles/mL for MIT9312 and MIT9313 cultures, respectively. Vesicle samples were resuspended in ~1 mL of PBS buffer and stored at -80 °C until extraction. Cell densities were measured by flow cytometry, corresponding to $1.29 \times 10^8$ cells/mL and $3.78 \times 10^7$ cells/mL for MIT9312 and MIT9313 cultures, respectively. Following vesicle isolation, cell biomass from each strain was collected by filtering approximately 150 mL onto three separate Durapore (0.22 µm) filters. Total cells on each filter

(~150 mL per filter) were $1.94 \times 10^{10}$ cells and $5.67 \times 10^{9}$ cells for MIT9312 and MIT9313, respectively. Artificial seawater media (AMP1) blanks were also collected on filters to serve as controls. Filters were stored at -80 °C until extraction.

Filters from the AMP1 media blank, MIT9312 cultures, or MIT9313 cultures were thawed, cut using sterile scissors and filters were resuspended (separately) in 1 mL of 50 mM ammonium bicarbonate (AMBIC) with 5 mM ethylenediaminetetraacetic acid disodium salt dihydrate (EDTA) buffer (pH 8) in a microcentrifuge tube containing glass beads. Four vesicle samples were extracted, corresponding to the PBS buffer blank, AMP1 media blank, MIT9312 vesicles and MIT9313 vesicles. Each vesicle sample (1 mL) was thawed, additionally suspended to a final concentration of 50 mM AMBIC, 5 mM EDTA (pH 8) and then transferred to a microcentrifuge tube containing glass beads. Each filter or vesicle suspension was subjected to bead-beating for 30 seconds at 6 m/s, frozen at -20 °C for 45 minutes, and thawed. The bead-beating and freeze-thaw cycles were repeated twice followed by a third bead-beating cycle and an overnight freeze at -20 °C. Following the overnight freeze, filter resuspensions were thawed and lysed cells with beads were vortexed and then centrifuged at 7,000 rpm (4,602 rcf) for 2 minutes. A 500 µL aliquot from each tube containing the corresponding filter was used for downstream proteomic processing (see below). For the purposes of this experiment, bulk protein concentration for lysed cell extracts were estimated from cell density measurements. For downstream proteomics analyses, we assume an 80% extraction efficiency to lyse cells or vesicle contents by bead-beating. Thus, we estimated for MIT9312 cells and MIT9313 cells that approximately $3.87 \times 10^{9}$ cells or $2.27 \times 10^{9}$ cells, respectively, were extracted and used for downstream proteomics processing. For vesicle extracts, we estimate that approximately $8.9 \times 10^{11}$ and $5.8 \times 10^{11}$ particles for MIT9312 and MIT9313 culture, respectively, were used for downstream proteomic processing. Due to limited vesicle biomass, vesicle samples were prepared to be processed *only* by trypins/Lys-C proteolytic digestion. Each 20L batch of *Prochlorococcus* culture yielded only enough biomass for one proteomics analysis.

Proteins from lysed cells or vesicles were subjected to in-solution proteolytic digestion following identical procedures and accounting for differences in cell/particle counts and sample volumes. First, RapiGest SF (Waters), an acid labile surfactant, was added to help facilitate protein solubilization (0.06% *w*/*v*). Next, protein extracts from lysed cells or vesicles underwent disulfide reduction with tris(2-carboxyethyl)phosphine (TCEP; to a final concentration of 10 mM) for 1 hour at room temperature in the dark. Reduced samples were then alkylated with iodoacetamide (IAM; to a final concentration of 30 mM) for 1 hour at room temperature in the dark. Excess IAM was quenched by the addition of dithiolthreitol (DTT; to final concentration of

5 mM) for 1 hour at room temperature in the dark. Each protein extract was proteolytically digested with mass spectrometry-grade trypsin/lys-C mix (Promega) at an *estimated* substrate-to-enzyme ratio of 25 (*w*/*w*; based on cell density conversion to protein concentration) for 17 hours at 37 °C containing a final concentration of 0.05% *w*/*v* RapiGest, 50 mM AMBIC and 5 mM EDTA (pH 8). Additionally, to generate more comprehensive cellular proteomes, a subset of samples from *only* the cellular fraction from MIT9312 and MIT9313 were also proteolytically digested with mass spectrometry-grade Glu-C (Promega) at an *estimated* substrate-to-enzyme ratio of 25 for 17 hours at 37 °C containing a final concentration of 0.05% *w*/*v* RapiGest, 50 mM AMBIC and 5 mM EDTA (pH 8).

Following protease digestion, RapiGest was hydrolyzed using established manufacturer protocols by the addition of trifluoroacetic acid (0.5% final *v*/*v*, pH < 2) that concurrently terminates protease activity, heated for 45 minutes at 37 °C and centrifuged at 15,000 rpm (21,130 rcf) at 4 °C for 15 minutes to precipitate the water immiscible decomposition product, dodeca-2-one. The supernatant was removed and dried to near-dryness using a centrivap concentrator (Thermo Savant SPD2010 SpeedVac). Digested samples were desalted using a MacroSpin C18 column (NestGroup) following an established manufacturer protocol. Samples were eluted from the MacroSpin columns using two times 200 µL of 80% LCMS-grade acetonitrile with 20% LCMS-grade water containing 0.1% LCMS-grade formic acid. Desalted samples were concentrated using the centrivap to near dryness. All samples including media controls were resuspended in 95% LCMS-grade water with 5% LCMS-grade acetonitrile containing 0.1% LCMS-grade formic acid. The resuspension solution also contained an internal standard of synthetic peptides (Hi3 *Escherichia coli* Standard, Waters) at 50 fmol/µL. The resuspension of cellular protein samples was to an approximate $5.5 \times 10^7$ cells/µL. Due to limited biomass, vesicle samples were resuspended in the smallest possible volume of 9.5 µL to an approximate $9.4 \times 10^{10}$ particles/µL and $6.2 \times 10^{10}$ particles/µL for MIT9312 and MIT9313 cultures, respectively. Media and control samples (AMP1 and PBS buffers) were resuspended identically to corresponding cellular or vesicle samples. Prior to injection and analysis by LCMS, iRT retention time standard (Biognosys) was spiked at a ratio of 1:20 iRT to sample.

Samples were injected onto the column of a Waters ACQUITY M-class UPLC with an injection volume of 2 µL, corresponding to on-column estimates for MIT9312 cultures of $1.05 \times 10^8$ cells and $1.78 \times 10^{11}$ vesicles, and for MIT9313 cultures of $1.08 \times 10^8$ cells and $1.17 \times 10^{11}$ vesicles. (Note these on-column values will be used in comparative protein analyses as normalization for a 'per cell' or 'per vesicle' basis, see below). Triplicate injections were conducted for cellular samples and due to limited biomass and sample volume, *only* one injection for vesicles samples. Peptide separation was performed by reversed-phase chromatography using a nanoACQUITY HSS T3 C18 column (1.8 µm, 75 µm x 250 mm; 45 °C) with an ACQUITY UPLC M-class Symmetry C18 trapping column (180 µm x 20 mm). The peptides were trapped at a flow rate of 5 µL/min at 99% A for 3 minutes. A flow rate of 0.3 µL/min was used over a gradient between LCMS-grade water (A) and LCMS-grade acetonitrile (B), both modified with 0.1% LCMS-grade formic acid. The total 145 minute gradient method for the separation of the peptides started at 95% A and ramped to 60% A over the course of 120 minutes. The gradient then switched to 15% A at 122 -127 minutes followed by a ramp back to starting conditions 95% A at 128-145 minutes.

The M-class UPLC was coupled to a Thermo QExactive HF Orbitrap high-resolution mass spectrometer (HRMS) equipped with a nano-electrospray (NSI) source made in-house following the University of Washington Proteomic Resource (UWPR) design. Using a MicroTee (PEEK, 0.025” OD), the commercial analytical column was connected to a commercial emitter (PicoTip, Waters) and the liquid path was applied a high voltage through a platinum wire (adapted from UWPR design). All analyses were carried out in positive mode at a NSI spray voltage of 2.0 kV. Data was collected using data dependent acquisition using Xcalibur 4.0 data acquisition software (Thermo Fisher). The $MS^1$ scan range was 400-2000 *m/z* at 60,000 resolution with a maximum injection time of 30 ms and automated gain control of 1e6. Following each $MS^1$ scan, data-dependent $MS^2$ (dd-$MS^2$) was set to perform on the top 10 ions in a data-dependent manner at 15,000 resolution with a normalized collision energy of 27 eV. Additional selection criteria for dd-MS2 were as follows: maximum injection time of 50 ms with an automated gain control of 5e4, the isolation window was 1.5 Da and dynamic exclusion was set at 20 sec.

Data processing was conducted using the software from the trans-proteomic pipeline (TPP v.5.1.0) (Nesvizhskii *et al.*, 2007). Briefly, raw data was converted to mzML and searched using COMET (Eng *et al.*, 2013) against a FASTA protein database consisting of either MIT9313 (Uniprot accession PROMM, accessed September 2015) or MIT9312 (Uniprot accession PROMP, accessed March 2017), Hi3 standard (*E.coli* chaperone protein; P63284), iRT retention time standard, and a concatenated set of randomized sequences. We evaluated combinations of additional COMET search parameters and for the purposes of this manuscript, COMET parameters (designated COMET*a*) included no enzyme specificity, carbamidomethylation of cysteine residues as a fixed modification (+57.0215 Da), and oxidation of methionine residues (+15.9949 Da) and clipping of N-terminal methionine as variable modifications. To aide in identifying intracellular prochlorosin precursors, a second set of COMET searches (herein referred to as COMET*b* search) was also performed with these same parameters using only the no enzyme specificity and with all variable modifications, including clipping of N-terminal methionine, oxidation of methionine residues (+15.9949 Da), carbamidomethylation of cysteine residues (+57.021464 Da), and dehydration of serine and threonine residues (-18.010565 Da) to 2,3-didehydroalanine (Dha) and (*Z*)-2,3-didehydrobutyrine (Dhb), respectively. Files were then searched in PeptideProphet with greater than 90% peptide probability and further analyzed using iProphet and ProteinProphet within the TPP. The list of protein identifications was filtered using a 95% or greater protein probability as calculated from iProphet corresponding to a FDR < 1%. To ensure adequate comparison of relative protein abundances across cellular and vesicle proteomes of *Prochlorococcus* strains MIT9312 and MIT9313, we opted to further filter the identified protein datasets where at least one peptide (or spectral counts) must be observed consistently across individual injections and the protein probability within each replicate must be greater than 95%. Abacus software (Fermin *et al.*, 2011) was used to help join and organize PeptideProphet and ProteinProphet data across replicates and treatment conditions.

Notably, we must consider here the differences in cell and vesicle biomass injected on-column for the two *Prochlorococcus* strains MIT9312 and MIT9313. Overall, vesicle samples had much lower biomass on-column and thus many peptides observed were closer to the limit of detection in HRMS. In DDA, the probability of a protein being identified is directly related to the abundance of a protein, which is made up of these observed peptide abundances. It is possible that more proteins are expressed in the vesicle samples of these *Prochlorococcus* strains; however, they were not detected because their peptide abundances exceeded our limit of detection. Overall, this lends to the lower percentage of the vesicle proteome being detected and more importantly, to the lower percent protein coverage observed. For instance, protein analyses in the vesicle proteomes resulted in a higher number of proteins identified based on only one or two peptide identifications (Table S1).

For label-free quantitative proteomics, the relative abundances of proteins across cells and vesicles are compared using normalized spectral counts, which were computed using the following normalization procedures. All raw spectral counts were first averaged across technical replicates and standard deviations of these averages were calculated for error (Table S1). Averaged spectral counts of proteins were normalized to the Hi3 internal standard to account for differences in NSI ionization of peptides due to sample matrix effects and then normalized on "per cell" or "per vesicle" basis by dividing by the amount of biomass injected on-column (see on-column values above; Shah *et al.*, 2019) (Table S2), resulting in a semi-quantitative estimate of 'normalized spectral counts per cell' or 'normalized spectral counts per vesicle'. To compare protein enrichments within *Prochlorococcus* cell and vesicle fractions, "per cell" or "per vesicle" spectral counts were then normalized on the basis of size, or biovolume, differences between a cell and a vesicle, herein referred to as "biovolume-normalized protein spectral counts" (Table S2). Estimated volume values were computed using an average radius of 375 nm for cells or 50 nm for vesicles (Biller *et al.*, 2014), corresponding to biovolumes of 0.221 $\mu m^3$ and 0.000524 $\mu m^3$, respectively. Normalized spectral counts "per cell" or "per vesicle" were then divided by these volumes to yield "normalized spectral counts per cell biovolume ($\mu m^{-3}$)" (with propagated error) and "normalized spectral counts per vesicle biovolume ($\mu m^{-3}$)" (Table S2). Subcellular localization assignments are based on a combination of predictions from the Uniprot database (https://www.uniprot.org), results from the PSORTb algorithm (V3.0.2) (Yu *et al.*, 2010), amended with data from TMHMM 2.0 (Krogh *et al.*, 2001) and SignalP (V4.1) (Nielsen, 2017), based on the Gram-negative bacteria model.

*Metabolomics*

Cell pellets and extracellular vesicles originating from triplicate 20 L batch cultures of *Prochlorococcus* strains MIT9312 and MIT9313 were extracted using a modified Bligh and Dyer technique with cold 1:1 methanol/water (aqueous phase) and cold dichloromethane (organic phase) (Bligh and Dyer, 1959; Boysen *et al.*, 2018). Cell pellets were manually disrupted by bead beating during the extraction, as described by (Boysen *et al.*, 2018). 1.7 x $10^{10}$ – 2.3 x $10^{10}$ cells were extracted per replicate for MIT9312 cells, and 3.1 x $10^9$ – 9 x $10^9$ cells for MIT9313. Vesicles were extracted without bead beating, with each replicate containing 4.9 x $10^{10}$ – 7.1 x $10^{10}$ vesicles for MIT9312 and 3.8 x $10^{11}$ – 6.2 x $10^{11}$ vesicles for MIT9313. Process blanks (MilliQ water), media blanks, and PBS (vesicle suspension buffer) were extracted and analyzed alongside each sample set. Authentic standards of all targeted metabolites were analyzed within sample batches to establish retention times of each standard. Isotope-labeled internal standards were added to samples before and after extraction to aid in normalization, exactly as described in Boysen *et al.*, 2018. Pooled-samples were run several times throughout the sample batch to track instrument performance and aid in normalization (Boysen *et al.*, 2018).

Metabolite separations were achieved using reversed-phase (RP, for aqueous and organic extracts) and hydrophilic interaction liquid chromatography (HILIC, for aqueous extracts only), as detailed in Table S4 and (Boysen *et al.*, 2018). Data were collected in positive ion mode for RP analyses and in positive and negative mode (using polarity switching) for HILIC analyses. For every sample, data were processed in four subgroups, defined by phase, chromatography, and ion mode: RP-organic-positive (RPOrgPos), RP-aqueous-positive (RPAqPos), HILIC-aqueous-positive (HILICAqPos), HILIC-aqueous-negative (HILICAqNeg).

After extraction, samples were run within 24 hours for HILIC, and 96 hours for RP. Sample extracts were stored at -80°C between extraction and analysis. Metabolite data were collected using a Thermo QExactive HF with collection parameters reported in (Boysen *et al.*, 2018). For untargeted analysis, we collected fragmentation spectra ($MS^2$) on all samples, using the same MS parameters as previously described (Heal *et al.*, 2019). Pooled samples were run several times throughout the sample batch to track instrument performance and aid in normalization to minimize obscuring variability that is inherent to LC-MS analysis. This normalization process is described in detail in (Boysen *et al.*, 2018).

For targeted data, individual metabolite features were integrated using Skyline Daily (MacLean *et al.*, 2010) and subjected to an in-house quality control protocol (Boysen *et al.*, 2018) that rejects features with inappropriate retention times or poor mass matches. Quality controlled data were normalized by the Best-Matched Internal Standard (B-MIS) normalization protocol (Boysen *et al.*, 2018). Blank values were subtracted from normalized data; blank corrected data were normalized for biological variability, based on biovolume (see above).

Untargeted data files were converted from Thermo.RAW file format to the universal .mzXML format, using MSConvert software (Chambers *et al.*, 2012). Untargeted data were further processed using MS-DIAL software (Tsugawa *et al.*, 2015) using parameters reported in Table S9. MS-DIAL was used to pick, align, and integrate mass features from raw datasets. Mass features whose area were not at least ten times the average peak area of relevant analytical blanks were excluded from downstream analysis. In addition, mass features that were likely $^{13}C$, $^{15}N$, or $^{34}S$ isotopologues of other mass features were identified and also excluded from further analysis. Aligned, integrated, and filtered mass features were normalized to account for non-biological variability using B-MIS (Boysen *et al.*, 2018). Blank corrected data were normalized for biological variability based on biovolume.

Identification of mass features within untargeted data sets was accomplished via dereplication of the mass feature list against several databases, yielding identifications of variable confidence. We ranked confidence in mass feature identifications according to existing literature (Sumner *et al.*, 2007). All mass features were searched against an Ingalls Lab in-house database of authenticated standards (https://github.com/IngallsLabUW/Ingalls_Standards), LOBSTAHs output, and MassBank (Horai *et al.*, 2010). Searches against the Ingalls Lab database were performed by comparison of exact *m/z* and RT between mass features and authentic standards. Matches to the in-house database yield the highest confidence identifications (confidence level 1) since authentic standards are available for each of these compounds.

Matches to the LOBSTAHs database were performed by comparison of parent *m/z* and manual inspection of $MS^2$ spectra and these mass features were not further considered. Matches to MassBank were established by searching for compounds with a matching parent *m/z* and $MS^2$ spectra with cosine > 0.8 using algorithms used in previous literature (Horai *et al.*, 2010). MassBank identifications were verified by manual inspection of $MS^2$ spectra. In most cases, identifications made on the basis of $MS^2$ spectra, in the absence of an authentic standard, yielded a confidence level 2 identification, indicating the putative assignment of a compound identification. In a small number of cases, these yielded a confidence level 3a identification, indicating the mass feature could be putatively assigned to a compound class, but not a specific member of that class.

The mass features with the largest MS signal which were not identified by comparison to the preceding databases were manually searched in Metlin (Smith *et al.*, 2005) and the Global Natural Products Social Molecular Networking (GNPS) (Wang *et al.*, 2016) databases. Metlin searches were performed by initially screening for compounds (as proton adducts, $M+H^+$) with parent masses within 5 ppm of each *m/z*. Putative compound identifications were assigned after performing visual comparison of $MS^2$ spectra. Matches to the GNPS were established by searching for compounds with a matching parent mass within 5 ppm of each *m/z* and $MS^2$ spectra with cosine > 0.8. Results from GNPS searches were verified by manual inspection of $MS^2$ spectra. Similar to putative compound assignments generated by MassBank searches, assignments derived from Metlin and GNPS were also considered confidence level 2 matches.

The top 1000 largest peaks in each vesicle sample (considered by area) were further subjected to molecular networking analysis. Cosine similarity scores were calculated in a pairwise fashion for all qualifying mass features using the same algorithm as our database searches (Horai *et al.*, 2010). Features with cosine scores between 0.6 and 0.95 were connected by nodes in a network plot. Network plots were created using R's ggnetwork package (Briatte, 2020) applying the Fruchter Manreingold layout (Fruchterman and Reingold, 1991). Nodes can be overlaid with metadata, including identifications and associated confidence levels, enabling putative structures of unknown, but structurally-related metabolites, to be proposed with a degree of confidence. In this study, molecular network plots were used to support the assignment of hypothetical compound identifications made in the absence of authentic standards or published reference spectra. Putative identifications assigned using information from molecular network plots was considered level 3b, as the molecular network plots confirmed a set of mass features were structurally related and likely to be members of a class of compounds.

*Vesicle labeling and interaction assays*

*Alteromonas* and *Prochlorococcus* vesicles were labelled with an Alexa Fluor 488 5-SDP ester dye (Molecular Probes/ThermoScientific). 200 µl of vesicles were mixed with 5 µl of dye in DMSO (1 mg/mL) in fresh 0.1 M sodium bicarbonate buffer (pH 8.3) and incubated for 1 hour in the dark at room temperature with gentle mixing. Excess unbound dye was removed by washing the sample 3x with 1x PBS in an ultracentrifuge (45 minutes / 100,000 x*g* spin). A control labeling reaction was also carried out using just the PBS buffer. Final concentrations of labeled vesicles were determined using the NanoSight. We chose this covalent amine labeling approach, as opposed to the use of perhaps more general vesicle labeling approaches using lipophilic dyes such as FM4-64, for multiple reasons. First, we have found that lipophilic dyes tend to form micelles on their own in seawater which are difficult to separate from vesicles, adding a large background signal to the analysis. Second, we had concerns about the stability of lipophilic membrane labeling in this context, as dye could potentially leave vesicle membranes and transfer to cells absent a direct vesicle-cell interaction. DNA staining would be expected to only target a tiny fraction of vesicles with sufficient DNA content (Biller *et al.*, 2017), whereas amine labeling, presumably targeting primarily surface proteins, should allow a larger fraction of all vesicles to be tracked.

To examine the ability of vesicles to associate with representative phylogenetically distinct marine microbes, *Prochlorococcus* MIT9312 and MIT9313 (in Pro99 media), *Alteromonas* MIT1002 and *Thalassospira* MIT1004 (Biller *et al.*, 2017) (in ProMM media), and *Polaribacter* MED152 (in Pro99 media, supplemented with 5 g/L peptone and 1g/L yeast extract) were all grown to mid-exponential growth phase at 24 °C. *Pelagibacter* HTCC7211 was grown in AMS1 media at 22 °C. *Alteromonas* MIT1002 and *Thalassospira* MIT1004 are available from the authors upon request.

Approximately $10^9$-$10^{10}$ vesicles (labeled or unlabeled), or an equivalent volume of the PBS Alexa 488 labeling control, were added to 2 mL of culture at ~$10^5$ cells $mL^{-1}$ (final ratio: ~1000:1 vesicles:cells). Cultures and vesicles were incubated for 2 hours at the normal growth temperatures. 0.5 µm blue-excited reference beads were added to each sample, and data was collected on an Influx flow cytometer (Cytopeia/BD, Franklin Lakes, NJ). Cells were excited using a blue 488 nm laser and monitored for chlorophyll (692/40 nm emission) and Alexa 488/GFP (530/40 nm emission) fluorescence. Flow cytometry data was collected for a set amount of time for each strain tested, with >25,000 cellular gate events collected per sample. For washing controls (Fig. S2), 1 mL aliquots of cell-vesicle mixtures were twice pelleted at 7500 x*g* for 5 minutes and washed with fresh media prior to flow cytometry.

All flow cytometry data was analyzed using FlowJo (V10.5). *Prochlorococcus* cell populations were gated based on forward scatter and 692/40 nm chlorophyll signatures; heterotrophs were gated based on forward scatter and 530/40 nm fluorescence. Representative gates are shown in Figs. S8/S9. Median 530/40 nm cellular fluorescence values were calculated from these population gates and normalized to the fluorescence of the internal reference beads. Statistical comparisons to evaluate the shift in 530/40 nm fluorescence were computed using the nonparametric Chi Squared T(x) statistic. Per the FlowJo documentation, we first determined a biologically relevant minimum value of T(x) by comparing the cellular populations when incubated with either PBS buffer alone vs with non-fluorescently labeled vesicles to assess the background variation. Vesicle-cell interactions were only considered biologically significant if the Chi Squared T(x) comparison between cells incubated with labeled vs. unlabeled vesicles was at least 4x greater than the maximum T(x) value obtained among background comparisons, indicating a $p < 0.01$.

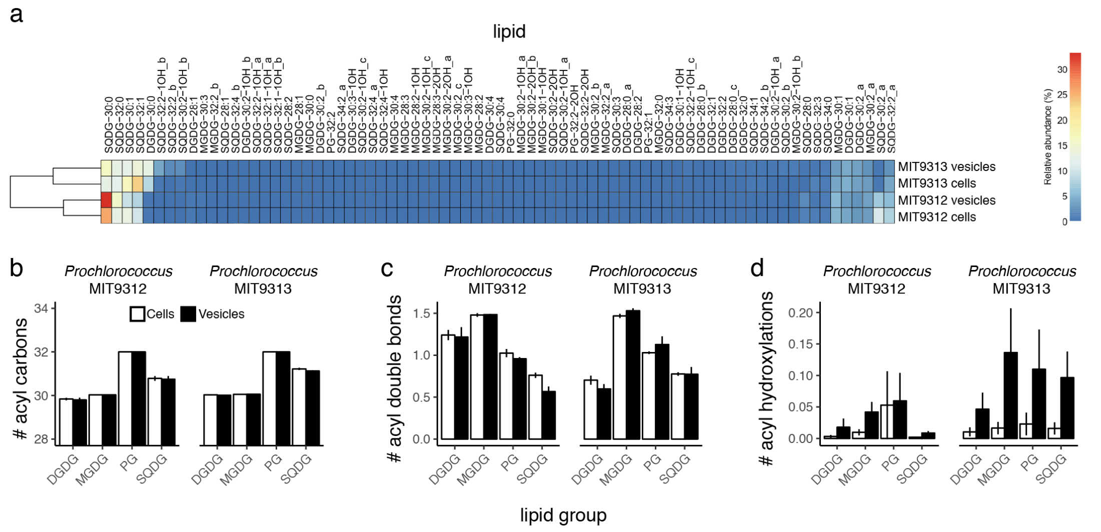


**Figure S1.** Comparative lipidomics of *Prochlorococcus* vesicles. (a) Heatmap and dendrogram showing relative distribution and hierarchical cluster analysis of lipid results. (b-d) Average number of (b) acyl carbons; (c) acyl double bonds; and (d) acyl hydroxylations of the total polar lipid pool in *Prochlorococcus* MIT9312 and MIT9313 cells and vesicles. Data indicate mean (+/- SD) of three biological replicates. MGDG, monoglycosyl diacylglycerol; DGDG, diglycosyl diacylglycerol; PG, phosphatidylglycerol; SQDG, sulfoquinovosyl diacylglycerol.

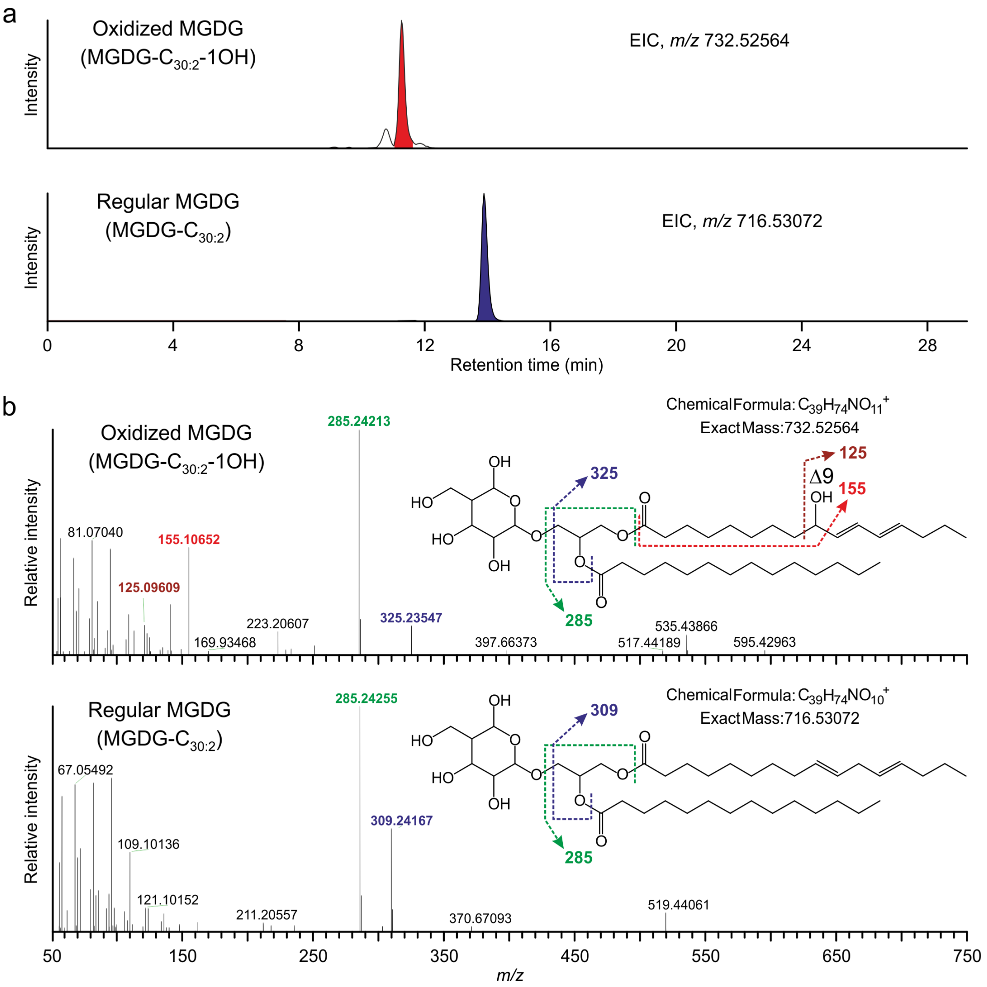


**Figure S2.** Identification of oxidized polar lipids. (a) Extracted ion chromatograms (EIC) of monohydroxylated ($[M+NH_4]^+$ at *m/z* 732.5) and regular MGDG-$C_{30:2}$ ($[M+NH_4]^+$ at *m/z* 716.5) in vesicles from *Prochlorococcus* MIT9313. (b) $MS^2$ spectra of ammoniated ($[M+NH_4]^+$) monohydroxylated MGDG-$C_{30:2}$ (MGDG-C30:2-1OH) and regular MGDG-$C_{30:2}$ in vesicles from *Prochlorococcus* MIT9313. MGDG, monoglycosyl diacylglycerol.

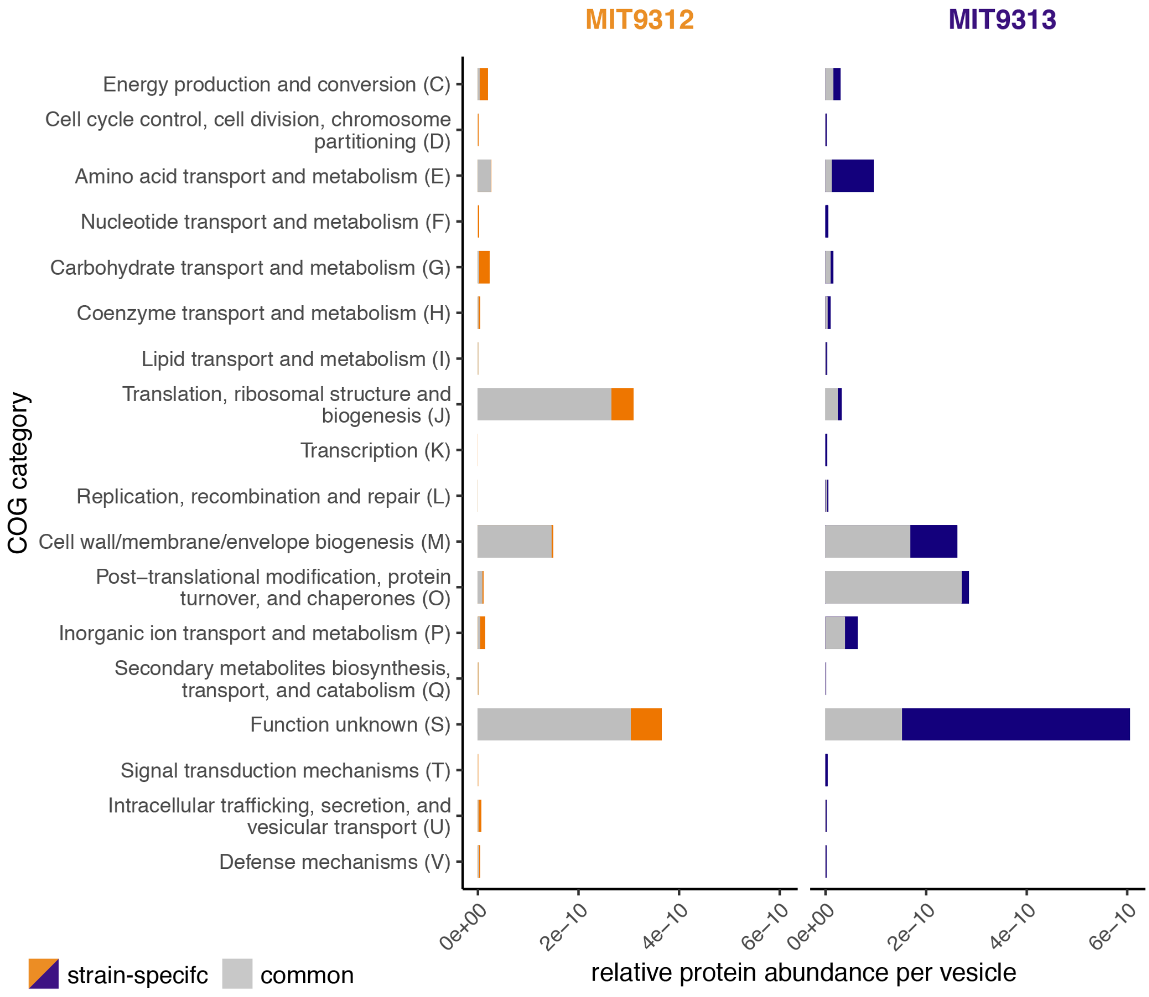


**Figure S3.** COG category distribution of proteins identified in *Prochlorococcus* extracellular vesicles. Values represent the relative abundance of proteins in the vesicle fraction (see also Dataset S2). Grey section indicates proteins found in both strains; colored regions represent proteins found uniquely in vesicles from each strain (MIT9312, orange; MIT9313, blue).

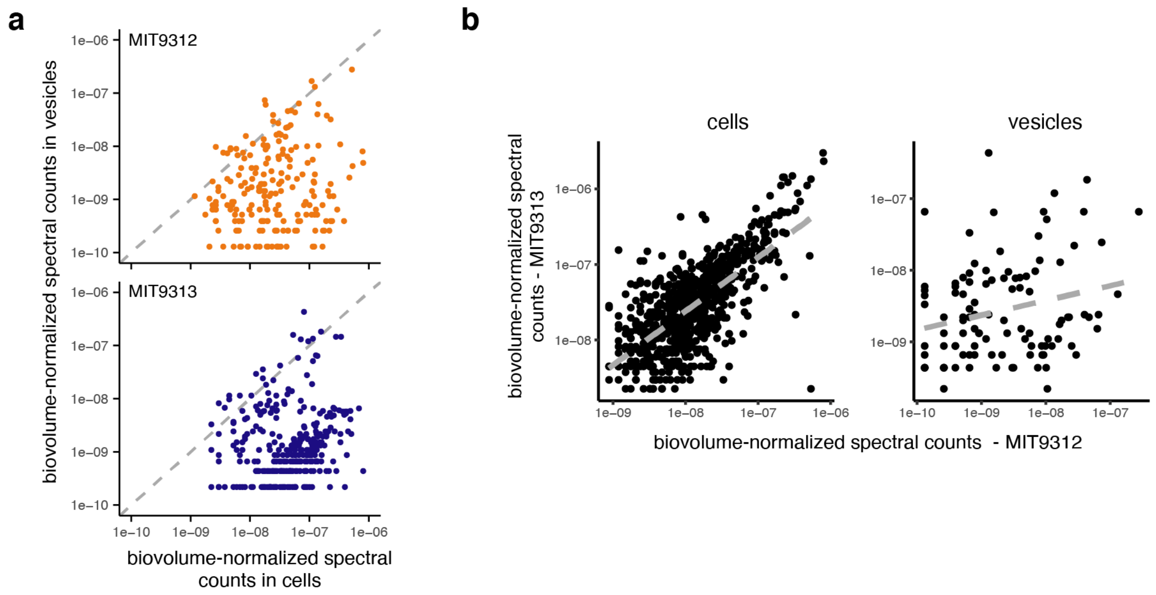


**Figure S4.** Comparative protein spectral counts. (a) Comparison of biovolume-normalized spectral counts for all proteins found in both the cells and vesicles of *Prochlorococcus* MIT9312 (above) and MIT9313 (below). Dashes represent the 1:1 line. (b) Comparison of biovolume-normalized protein spectral counts between the cell and vesicle fractions of the two *Prochlorococcus* strains. Dashed line represents best linear fit; $R^2$= 0.5 for cells, 0.057 for vesicles.

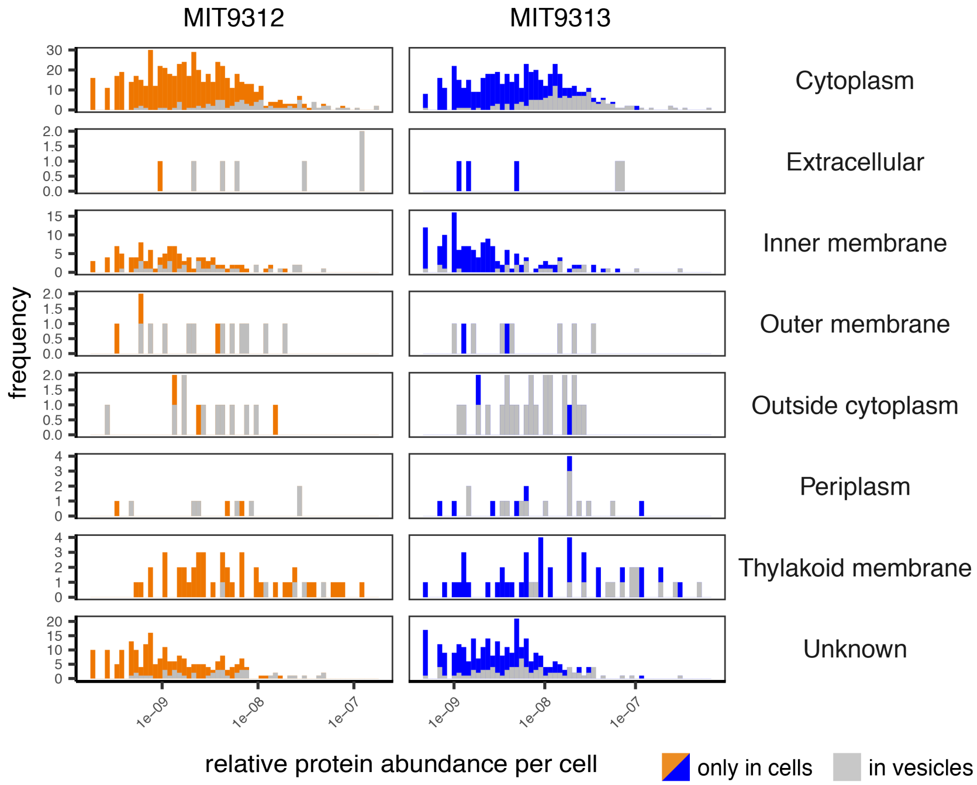


**Figure S5.** Vesicle packaging of proteins from different subcellular compartments. Histograms depict the relative protein abundance within MIT9312 (left) and MIT9313 (right) cells, separated by predicted subcellular compartment. Colored regions of the distribution indicate proteins found only in cells; grey regions indicate those cellular proteins also found exported within vesicles.

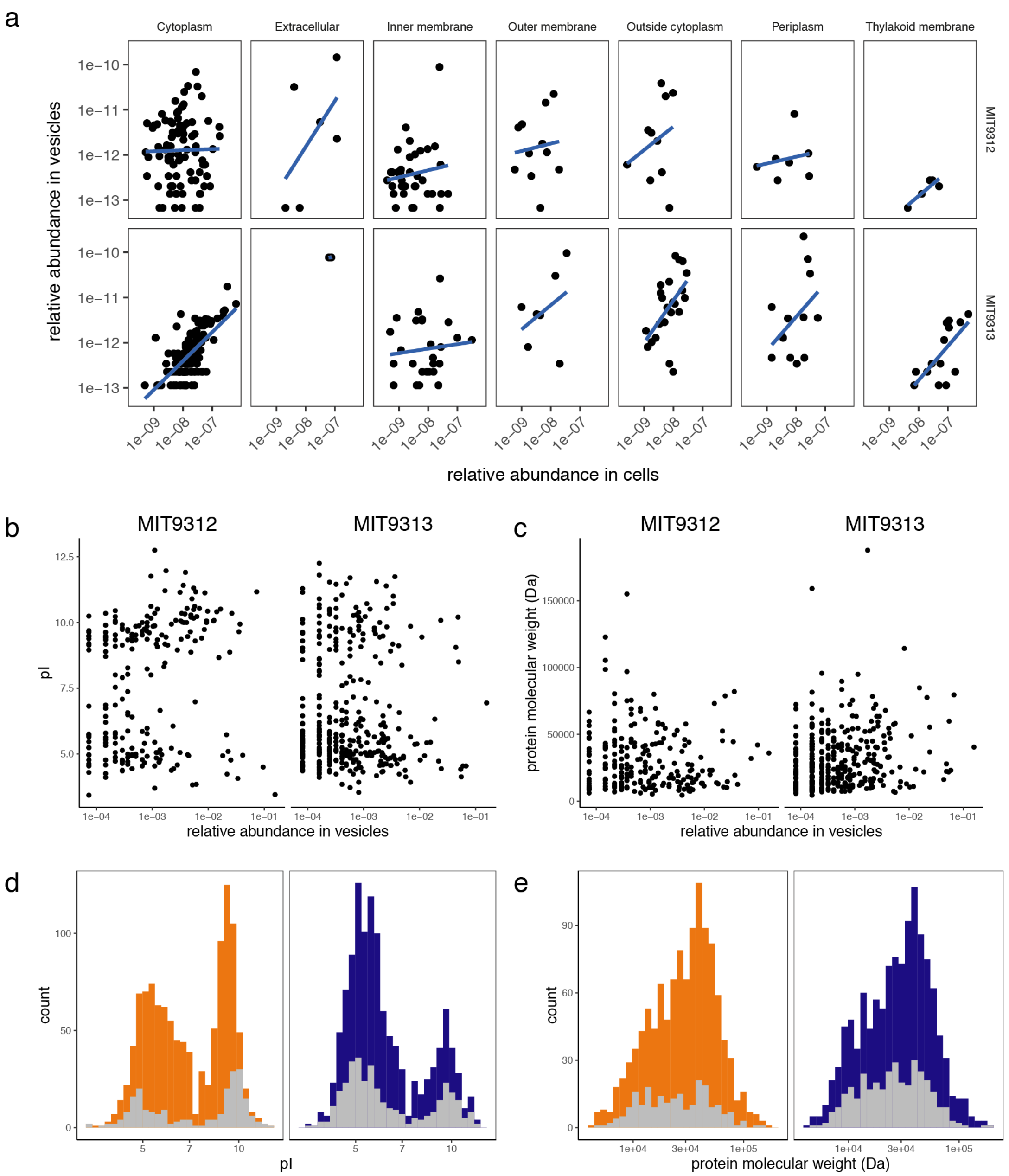


**Figure S6**. Vesicle proteome characterization. (a) Relationship between the abundance of proteins in the vesicle and cellular fractions, separated by the predicted subcellular localization of the protein within cells. Data are shown for strains MIT9312 (above) and MIT9313 (below); the line indicates the linear regression between the data for each panel. (b-c) Relationship between a protein's abundance in *Prochlorococcus* vesicles and its (b) calculated isoelectric point (pI) and (c) molecular weight. (d-e) Histograms showing the distribution of (d) pI values and (e) molecular weight of total predicted cellular proteome (color) vs those proteins identified within extracellular vesicles (grey). In b-e, MIT9312 is depicted on the left/orange; MIT9313 is on the right/blue.

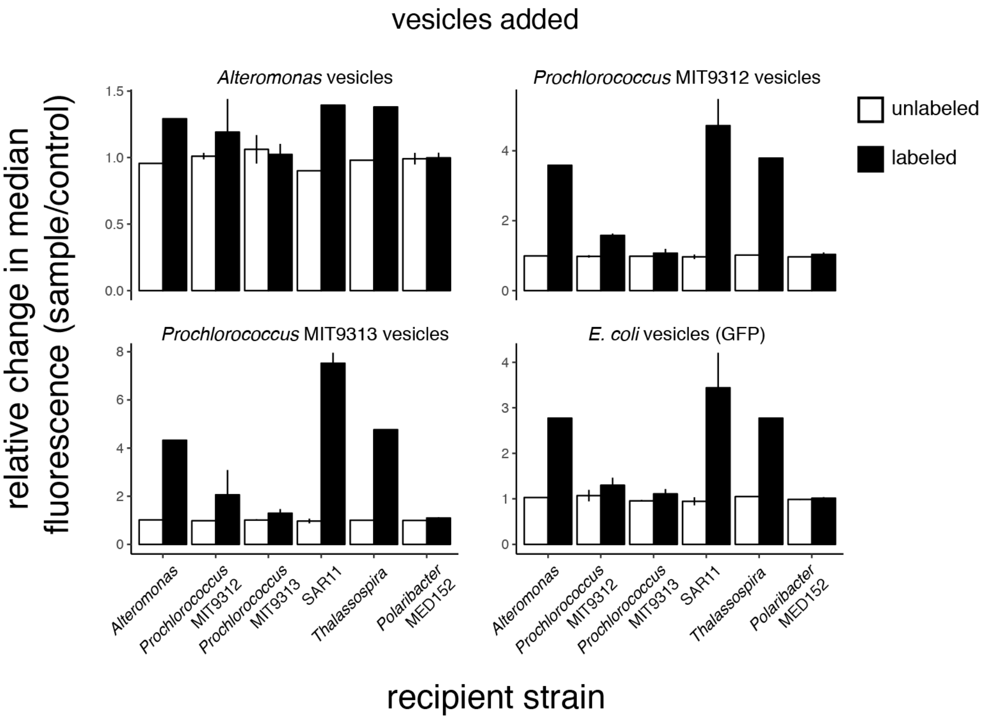


**Figure S7.** Relative shifts in median 530 nm fluorescence for six different marine microbes following incubation with vesicles from four different microbes. Fluorescence values were normalized against internal reference bead standards run within each sample. *Alteromonas* and *Prochlorococcus* vesicles were covalently labeled with an amine-reactive Alexa 488 dye; labeled *E. coli* vesicles were collected from a strain expressing a periplasm-localized GFP protein. Values indicate the normalized relative change in median cell fluorescence (+/- SD from *n*=2-3 biological replicates). Due to variation in cell sizes, growth rates, and fluorescence detection limits, quantitative comparisons of vesicle associations across both sources and recipients from these data are challenging and should not be over-interpreted.

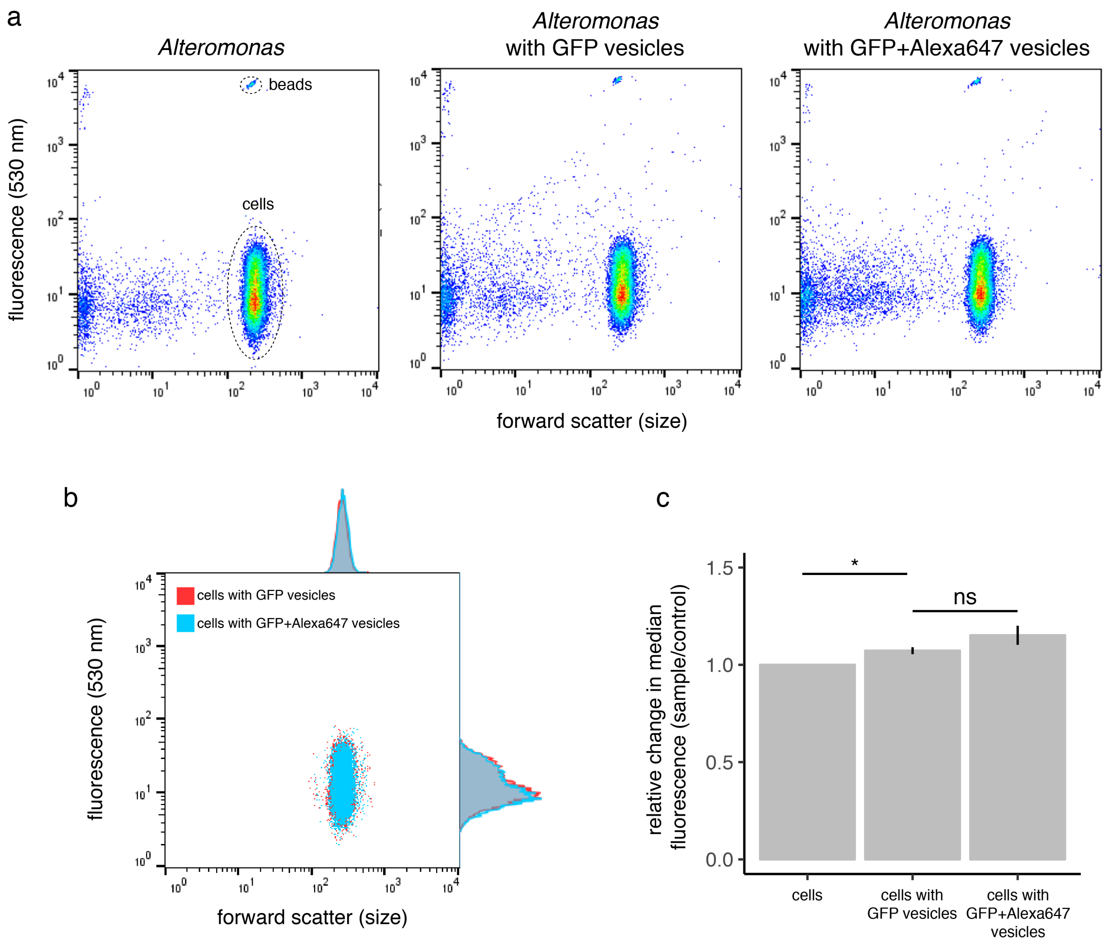


**Figure S8**. Characterizing the impact of surface labeling on vesicle-cell interactions. A sample of GFP-containing *E. coli* vesicles was labeled with an amine-reactive Alexa647 dye, which is structurally related to Alexa488 but with minimal excitation/emission in the 488/530 nm GFP channel. GFP and GFP+Alexa647 vesicles were added at equivalent concentrations, equal to those used for labeled *Prochlorococcus* vesicles, and incubated with heterotrophs as in Figs 1, S1. (a) Flow cytometry plots of *Alteromonas* cells vs cells incubated with the indicated vesicles. A representative example of population gating is shown by dotted lines. (b) Overlay dot plot and histograms comparing *Alteromonas* cell population fluorescence following vesicle incubation from part a. (c) Relative normalized shifts in median cell 530 nm fluorescence. While the median population fluorescence significantly increased upon incubation with GFP vesicles and GFP+Alexa647 vesicles (two-tailed $t$ test $p < 0.05$, and Chi Squared T(x) test $p \ll 0.01$, indicated by *), there was no significant difference between the response to the GFP vs GFP+Alexa647 labeled vesicles (two-tailed $t$ test $p=0.16$ and Chi Squared T(x) test; ns= not significant). The small average absolute difference between the response to GFP and GFP+Alexa647 vesicles (~7%) was below the magnitude of relative shifts considered significant in Fig. 4.

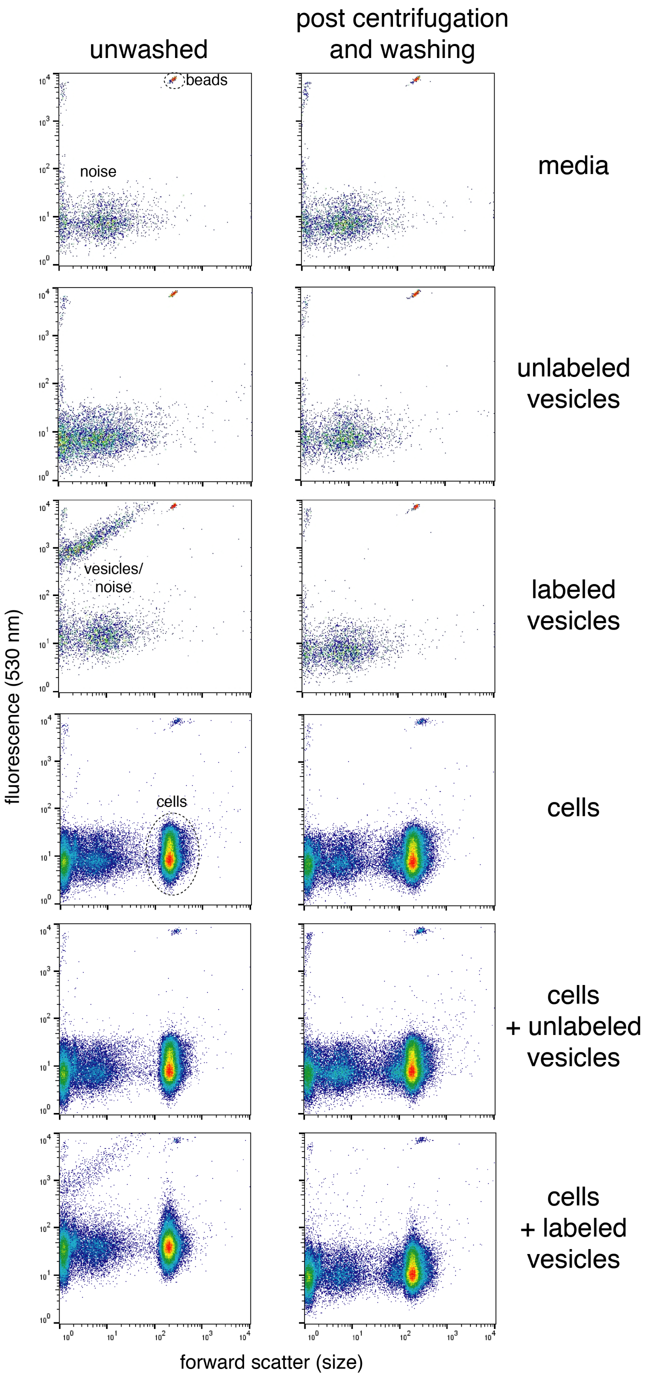


**Figure S9**. Stability of vesicle-cell interactions. Flow cytometry analysis of *Alteromonas* cells following incubation with a buffer control, unlabeled *Alteromonas* vesicles, or Alexa488-labeled *Alteromonas* vesicles. The 'unwashed' cells (left column) were run directly on the flow cytometer following incubation; the 'washed' population (right column) was assayed after being pelleted and washed twice with clean media (see Methods). Representative example of population gating is shown by dotted lines in the left column.
